# ‘Backpropagation and the brain’ realized in cortical error neuron microcircuits

**DOI:** 10.1101/2025.07.11.664263

**Authors:** Kevin Max, Ismael Jaras, Arno Granier, Katharina A. Wilmes, Mihai A. Petrovici

## Abstract

Neural responses to mismatches between expected and actual stimuli have been widely reported across different species. How does the brain use such error signals for learning? While global error signals can be useful, their ability to learn complex computation at the scale observed in the brain is lacking. In comparison, more local, neuron-specific error signals enable superior performance, but their computation and propagation remain unclear. Motivated by the breakthrough of deep learning, this has inspired the ‘backpropagation and the brain’ hypothesis, i.e. that the brain implements a form of the error backpropagation algorithm.

In this work, we introduce a biologically motivated, multi-area cortical microcircuit model, implementing error backpropagation under consideration of recent physiological evidence. We model populations of cortical pyramidal cells acting as representation and error neurons, with bio-plausible local and inter-area connectivity, guided by experimental observations of connectivity of the primate visual cortex. In our model, all information transfer is biologically motivated, inference and learning occur without phases, and network dynamics demonstrably approximate those of error backpropagation.

We show the capabilities of our model on a wide range of benchmarks, and compare to other models, such as dendritic hierarchical predictive coding. In particular, our model addresses shortcomings of other theories in terms of scalability to many cortical areas. Finally, we make concrete predictions, which differentiate it from other theories, and which can be tested in experiment.

## 1. Introduction

### 1.1. Background

Deep learning is the state of the art in training artificial neural networks (ANNs), with immense advances in practical applications made over the past decade. On the other hand, neuroscience has revealed a zoo of different brain areas, connection schemes and cell types. However, models taking into account such biological constraints have only produced limited understanding of how microscopic learning leads to complex function. Adapting individual components in a very complex network requires a meaningful local representation of errors; this has sparked the ‘backpropagation and the brain’ hypothesis [1, 2], i.e., that biological neural networks learn via a form of the error backpropagation algorithm (BP) powering deep learning.

However, the biological implausibility of BP has been a long-standing point of objection [1, 3]. Why would one then still consider backpropagation as a viable candidate for explaining learning in the brain? We argue that this is a consequence of applying Occam’s razor to physiological evidence: Error/mismatch encoding responses have been observed in a variety of brain areas and across species [4–12]. In particular, experimental findings suggest that cortical layer 2/3 (L2/3) pyramidal cells (PYR) function as error coding neurons [12–19]. Error neurons have also shown to emerge naturally in simulated RNNs trained under constraints of energy consumption [20]. If cortex has explicit error units, and all neurons are plastic, then it should arguably use such errors to their fullest.

Currently, BP appears to be the most efficient way for training multi-area networks in terms of generalizability, scalability, and task complexity. This is because BP propagates local errors across all areas, tailored to each neuron according to its contribution to task performance (credit assignment). Since BP works so well on artificial networks, our aim is thus to discover whether the cortical hierarchy could implement a bio-plausible variant of the BP algorithm. In this work, we suggest a concrete circuit implementation, which provides experimentally testable predictions and performs well on many different benchmarks, and compare it to alternative suggestions.

### 1.2. Related work

A contentious point of discussion is the (in-)compatibility of BP with the predictive processing framework (predictive coding, PC). However, as highlighted by [21–25], there is a fundamental relationship between the computations performed in hierarchical predictive coding (hPC) and ANNs trained with error backpropagation. Crucially, this means that the ‘backpropagation and the brain’ hypothesis and predictive coding are not mutually exclusive. Instead, they can be seen as closely related learning algorithms, which have traditionally been employed on different task domains: predictive coding networks (PCN) are commonly trained to predict stimuli, which is a generative task, while ANNs historically arose as solutions to classification problems. It is important to realize that the task domains are related by a simple exchange of stimuli and latent, and an inversion of the network hierarchy. Our model is built on this insight, and able to reflect it in its connectivity, with either a generative or classifier configuration (see Section 2).

While PCN can be successfully trained on various tasks and scale to many areas [26, 27], most studies do not detail how biology can implement the required computations. For example, the classical PC model by Rao and Ballard [28] is limited in its biological plausibility due to issues of ideal signal transport between neurons, weight copying, strictly hierarchical connectivity, and neuronal non-linearities (see Section 4.2).

Recent work aims to address these issues one by one [29–41]. Two models stand out in particular due to their biological plausibility: the dendritic cortical microcircuit model of Sacramento et al. [33], and the dendritic hPC model by Mikulasch et al. [25], both of which propose that errors are constructed on dendrites of pyramidal cells via local inhibition. This novel idea fits well into the theory of balanced networks and predictive coding, but assumes local interneurons *exactly* copying pyramidal neuron activity, which is unlikely given that interneurons integrate inputs from different cells, sometimes even non-linearly [42]. Moreover, models with dendritic error construction do not scale to more than two areas, as we will demonstrate using extensive simulations.

In contrast, we propose a model using recurrently connected prediction and error streams, loosely implementing the forward/backward pathways of error backpropagation. Similar connectivity is explored in attention-gated reinforcement learning (AGREL) and variants [34, 38, 43–45], but without solutions for the above-mentioned issues of bio-plausibility.

### 1.3. Summary of our model

Our model addresses a large number of biological constraints, while establishing a clear relationship to BP. We model two distinct subpopulations of cortical pyramidal cells in each area: one error and one representation/prediction population, corresponding to layer 2/3 and layer 5 neurons respectively. Connections between error and representation units in different areas are based on the functional connectome of the primate visual cortex [46] (see Fig. 1 b1). Thus, inter-area connectivity does not need to be strictly hierarchical, as opposed to most PC models in the literature. Furthermore, there is no biologically implausible one-to-one matching of prediction and error units within each area.

**Figure 1:**
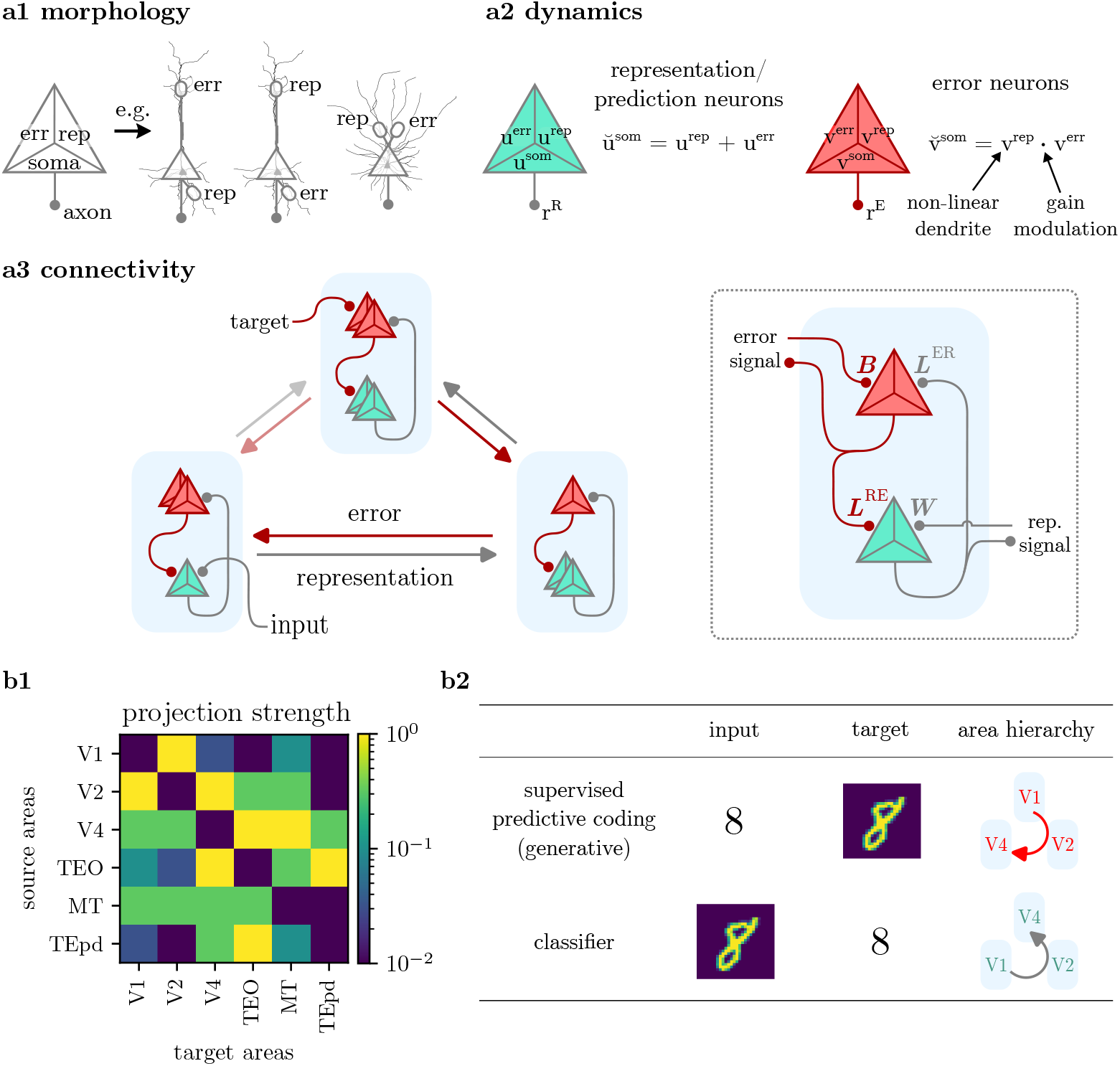
Neuron and network model. **a1)** Morphology: our model neurons have three compartments (soma, error and representation), which may be realized by various morphologies with apical, basal or perisomatic dendritic integration zones. **a2)** Dynamics: two types of pyramidal cells encode representations and errors, using distinct dynamics. **a3)** Connectivity: Between areas (left), two opposing streams of representation and error signals form, with preferential and weaker projections (opaque vs. semi-transparent arrows, respectively). Locally (right), representation units receive inputs from other areas onto their representation dendrite, and local error information on the error dendrite. Error units integrate afferent errors and local representations in their respective compartments. **b1)** Projection densities across the Macaque visual cortex, used for the inter-area connectivity in our model. Reproduced from Markov et al. 2014 [46]. **b2)** Generative and classifier configurations of the model differ by inputs, targets, and area labels. In the (supervised) predictive coding configuration, the model receives a latent representation (‘input’) and learns to predict bottom-up stimuli (‘target’). *Errors* are projected downstream in the hierarchy of cortical areas. The ANN configuration has input and target exchanged, the task becoming digit classification. The information flow is inverted, and *representations* are projected downstream.

In order to integrate both predictions and errors, each neuron is modeled with three separate integration zones (‘compartments’): one somatic and two separate dendritic compartments. For example, neocortical pyramidal cells are characteristically defined by their somatic, basal and apical dendritic integration zones [47, 48]. More generally, our model can be adapted to different neuronal morphologies, see Fig. 1 a1. Neuron dynamics are implemented in continuous time, without discrete phases of information transfer (i.e. no distinct forward/backward passes as in ANNs and some bio-plausible models [33, 34, 38, 43, 44]). The dynamics of each neuron type, in conjunction with its connectivity, encode their function as either error or representation units. Error units make use of non-linear dendritic dynamics, and communicate locally within each cortical column, providing a neuron-specific learning signal to each representation unit. This is important, because it treats error units as distinct cells with their own dynamics, without requiring knowledge of the somatic potential of local representation units (see Section 4.2). All neurons implement prospective rate coding [39, 40], modeling the diverse temporal signatures of neuronal outputs, and providing short-term memory to solve temporal tasks.

Learning is performed via a local Hebbian-type rule of the form presynaptic rate × postsynaptic voltage difference (delta rule). Similar to the voltage dynamics, learning is phaseless and always-on, as opposed to the vast majority of bio-plausible theories, such as PCN [21–24, 28], contrastive Hebbian learning [31, 49], the wake-sleep algorithm [50], difference target propagation and variants [30, 51, 52], AGREL [37, 38], and equilibrium propagation [32].

In this work, we will demonstrate the capabilities of our model to solve complex tasks and scale training to many cortical areas on par with ANNs, and compare in particular to the models of dendritic error construction [25, 33].

## 2. Theory

We now introduce the architecture of our model, and how it describes learning in a network with multiple areas. See Fig. 1 for a summary of the architecture. Basing the connectivity on that of the Macaque visual cortex [46], there is no strict hierarchy and connections between all areas are allowed (the equivalent of skip connections in ANNs). However, one may still describe a rough order between areas defined by the anatomical hierarchy of visual cortex [46, 53, 54] and the anterior temporal lobe [55] (Fig. 1 b1). The network admits two configurations in terms of this ordering (Fig. 1 b2): one where *errors* are passed from early to downstream areas (‘generative’ or ‘PC’ setup), and an inverted order, where *representations* are passed downstream (‘classifier’ setup). In this section, we keep the discussion of our model general to the configuration, and refer to the streams as representation (green in Fig. 1) and error streams (red) (note that in the context of predictive coding, representation neurons are also known as *prediction* units). Our model is thus quite general and can encompass various hypotheses of cortical architecture, such as different directions of information flow or projection densities between and within areas.

Stimuli and latent activations are presented to the ‘input’ and ‘target’ areas, which are defined depending on the configuration (Fig. 1 b2). For example, for the generative configuration, visual stimuli are presented to V1 as targets, while latent priors are provided to the downstream area V4 (‘input’); for sharp priors, e.g. in the form of labels, this amounts to supervised training. By modeling different latent priors, our model can also adapt other learning schemes, such as unsupervised learning as in the Rao-Ballard model (see Section 4.4). In each area, we model two populations of pyramidal cells, acting as representation and error units. Across areas, information is passed between representation units through unidirectional synapses *W*, while errors are calculated at the ‘target’ area, and projected between error units through uni-directional weights *B*. Within each area, representation and error units are connected via local weights *L*^ER^ (representation *→* error) and *L*^RE^ (error *→* representation). We model each pyramidal neuron with three dendritic integration zones: a somatic, a representation and an error compartment (Fig. 1 a1). Depending on the type of neuron (representation or error unit), these compartments perform different calculations and observe different connectivity (Fig. 1 a2).

Representation units integrate two streams of information: 1) representation units projecting from other areas onto a compartment with voltage *u*^rep^; and 2) same-area error units projecting locally onto an error compartment *u*^err^. Each representation unit *additively* integrates both streams in its somatic voltage *u*^som^:

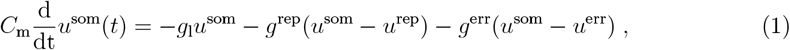

where *C*_m_ is the capacitance of the soma, *g*_l_ is the leak of the soma, and {*g*^rep^, *g*^err^} are the conductances of the two dendritic compartments. From these dynamics, one can see that representation neurons have an effective membrane time constant 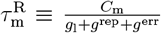. For simplicity, we set the leak potential *E*_l_ to zero for all neurons in our model; this can easily be included by adding a term *g*_l_*E*_l_ to all somatic dynamics.

In our model, the neuronal output is an instantaneous firing rate *r*(*t*). Across nearly all instances in the literature, *r*(*t*) is modeled as a function *φ* of the current membrane potential *u*(*t*) only (e.g., [24, 28, 33, 56]). However, this does not represent the temporal expressiveness of real neurons well, whose responses also depend on the dynamics of stimulating currents [57–59]. This effect can, to some degree, be captured by the ‘prospective (!) coding’ mechanism [39, 40, 59], where firing rates are functions of *u*(*t*) and instantaneous rate of change 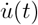. This means that the instantaneous firing rate is

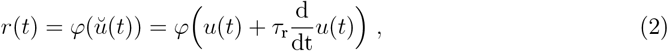

where *ŭ*(*t*) is defined as the *prospective* voltage with time constant *τ*_r_. For fast information transfer, neurons may have approximately equal prospective and membrane time constants, *τ*_r_ *≈ τ*_m_, such that the unavoidable lag introduced by the slow membrane is compensated by the prospectivity of the output. On the other hand, choosing smaller *τ*_r_ can implement neuron-wise short-term memory, enabling the network to solve temporal tasks (cf. Fig. 2). More generally, these time constants can be learned, enabling networks to adapt their memory to solve complex spatio-temporal tasks [39, 40]. For simplicity, we restrict ourselves to fixed *τ*_r_ in this work. We denote the prospective firing rate of representation and error units as *r*^R^(*t*) = *φ*(*ŭ*^som^) and 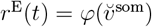 respectively. Note that we use linear activations for error neurons in our model (see below).

**Figure 2:**
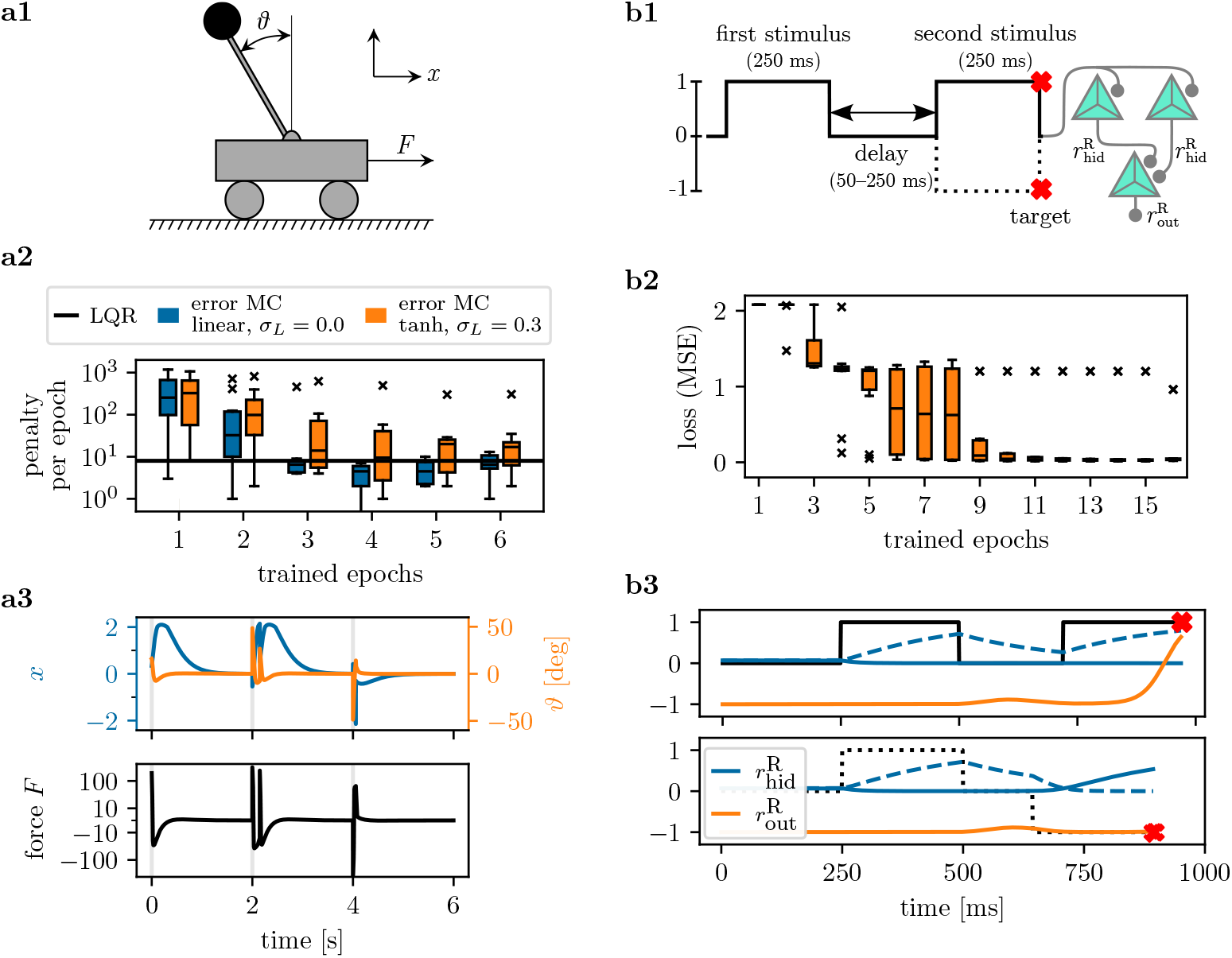
Error neuron microcircuits learn motor control and memory tasks. **a1)** Setup of cart pole task. The objective is to control the inverted pendulum such that *ϑ →* 0 and *x →* 0 are reached. **a2)** A microcircuit of size [4-1] successfully learns to stabilize the pendulum provided inputs 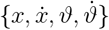. We show the number of failed runs for the linear-quadratic regulator (LQR) and two error neuron microcircuits with different activation functions and local noise level *σ*_*L*_. Shown are median and quartiles over 10 seeds. **a3)** Example trajectories for the trained microcircuit with tanh activation and *σ*_*L*_ = 0.3. The cart pole is reset every 2 s. **b1)** Delayed match-to-sample (DMS) task and network model. We show two of the four conditions with corresponding targets: ↑↑ (solid line) and ↑↓ (dotted). The time series is fed into a two-area network, each with [2-1] representation and error units (not shown). **b2)** Mean square error (MSE) during validation and testing; median and quartiles over 10 seeds. **b3)** Example dynamics for ↑↑ and ↑↓ after training, showing the representatsion neuron rates 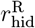 and 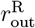. Note the different length of the delay for each of the two conditions. The network makes use of the slow membrane 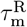 as short-term memory.

Error units encode a neuron-specific error signal in their soma, which they provide to local representation units. In order to encode useful errors, they need to integrate backpropagated errors from other areas with information from local representation units, in the form of the derivative of the activation 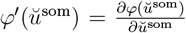. The integration of both signals is realized by two dendritic compartments in each error neuron: one receiving errors from other areas, and one for the reconstruction of *φ*^*′*^. The error backpropagation algorithm requires the multiplication of both signals – this may be realized by gain amplification of somatic activity by apical inputs, which has been observed in cortical L5 pyramidal cells [60] (see Discussion).

The soma of an error unit *v*^som^ therefore *multiplicatively* integrates its dendritic compartments,

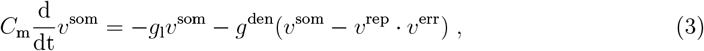

where *g*^den^ denotes the effective conductance coupling of both compartments to the soma. The modulation factor *φ*^*′*^(*ŭ*^som^) can be reconstructed in the dendrite using temporal noise *ξ*(*t*) propagating in the network, and the use of non-linear dendrites (see Methods, Section 6.4). For simplicity, we summarize the effective computation performed in both compartments as 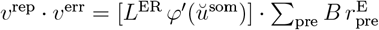.

Note that in our model, error signals are explicitly projected across areas, implementing the BP step *e*_*𝓁*_ = *φ*^*′*^(*u*_*𝓁*_)*⊙*[*W*_*𝓁*+1,*𝓁*_]^*T*^ e_*𝓁*+1_ (see Equation (5)). This differentiates our model from other PCN [21–25, 28], where errors are computed within each area as a local difference between top-down prediction and bottom-up stimulus. As we will demonstrate, this has benefits in terms of performance (Section 3.2) and biological plausibility (Section 4.2).

In this work, we do not address the handling of signed error signals. As neuronal outputs can only be non-negative, we need to explain how an error unit can communicate both positive and negative errors. This may be addressed by distinct populations for positive/negative errors, or transformed learning signals [61–65], see Section 4.4. For simplicity, we here assume linear activation functions for all error units, i.e. 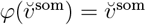.

We move on to the theory and learning rules for the weights, that is: *W* projecting representations, *B* projecting errors, and local weights *L*^RE^ and *L*^ER^. Note that there are no self-recurrent connections in our network, such that no population of representation or error neurons projects onto itself.

Representation-projecting weights *W* are the functionally relevant connections in the network, i.e., these weights are trained to minimize the cost/mismatch of a given task. To model the cortical connectome, we scale connectivity between different pairs of areas in *W* using the projection density of the Macaque visual cortex, see Fig. 1 b1. For learning of *W*, we use a form of the delta rule:

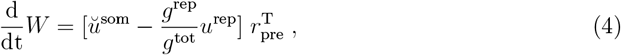

where 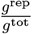 is a conductance-weighting factor, reflecting the coupling between soma and dendrite (see Methods). Note that the difference between soma and representation compartment encodes exactly the error signal required at the synapse, *ŭ*^som^ *− u*^rep^ = *u*^err^, for quasi-instantaneous neurons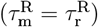.

Local weights are static in our model, and we assume that representation and error units are coupled by preferential targeting – i.e., there is only a loose one-to-one matching between both populations in each area. To model this, we initialize *L*^RE^ and *L*^ER^ as identity matrices with uniform noise, *L*^RE^ = ***1*** + *𝒰* (*−σ*_*L*_, *σ*_*L*_) and equally for *L*^ER^. In Section 3, we demonstrate how the network benefits from such loose matching (see also the Discussion).

Error-projecting weights: Exact backpropagation, like many realizations of predictive coding [21–23, 25], requires symmetric weights for the transport of representations and errors across areas, i.e. *B*_*𝓁,m*_ = [*W*_*m,𝓁*_]^*T*^. This is known as the *weight transport* problem, where two distant areas are required to share their synaptic weight information. Many solutions have been proposed in the literature, such as the Kolen-Pollack algorithm [35], Feedback Alignment [66] or Phaseless Alignment Learning (PAL) [67]. The latter harnesses intrinsic neuronal noise *ξ*(*t*) to train error-transporting weights *B*_*𝓁,m*_ to approximate [*W*_*m,𝓁*_]^*T*^, similarly to how we reconstruct *φ*^*′*^(*ŭ*^som^) in the error units. Furthermore, PAL is fully compatible with cortical microcircuit models, as demonstrated in [67]. Due to these two reasons, PAL is particularly suited to solve the weight transport problem in our model. It can be implemented by endowing the error-receiving synapses with the learning rule 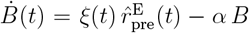, where *ξ*(*t*) is the noise on the potential of the post-synaptic error unit, 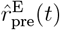 is the high-pass filtered, pre-synaptic rate of bottom-up error units, and *α* a scalar factor controlling homeostatic regularization. Note however that for computational feasibility, we here approximate PAL by setting error-projecting weights to a noisy transpose of representation weights (see Methods).

Finally, we stress that all information is processed simultaneously in our network (no phases), all information is locally available at the synapse, and learning is always-on.

## 3. Results

### 3.1. Motor control and spatio-temporal credit assignment

To demonstrate the capabilities of the error neuron microcircuit model for functional learning, we first train our implementation on two small benchmarks, see Fig. 2. The first is the cart pole task, a classical motor control problem also known as the inverted pendulum. The network receives four inputs: the current position and velocity of the cart 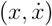, and angle and angular velocity of the pole 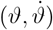. The aim is to produce a motor output (force *F*) which achieves an upright position of the pole (*ϑ* = 0) and navigates the cart to the position *x* = 0. The cart pole is initialized with *x* ∼ 𝒰 (−1, 1) and *θ* ∼ 𝒰 (−57^*°*^, +57^*°*^).

We train the networks to reproduce the motor control of a linear-quadratic regulator (LQR), an algorithm that optimises control for linear systems, and test performance using a simulation of the non-linear dynamics of the cart pole. Performance is measured by the penalty log, i.e. how often the pole falls or fails to reach the position *x* = 0. As Fig. 2 shows, error neuron microcircuits with five representation and error units each learn to solve the task, on par with the LQR. We demonstrate this both with the ‘ideal’ microcircuit, which has linear neuronal activation and perfect one-to-one matching between error and representation units, as well as the more realistic model, with non-linear *φ*(*ŭ*^som^) and noise *σ*_*L*_ on local weights.

Next, we show that error neuron microcircuits can solve tasks requiring memory. We use the delayed match-to-sample (DMS) task [68], equivalent to temporal XNOR. The task is to compare two sequential stimuli to identify if the stimuli match sign (*↑↑* or *↓↓*) or have opposing signs (*↑↓* or *↑↓*). The input is presented to the network as a one-dimensional time series, and therefore requires short-term memory to be solved. Task complexity is increased by an intermittent delay period with variable length.

In our model, short-term memory is encoded by the leaky integration of the membrane of representation units, and tunable memory length is achieved by the prospective coding mechanism (Section 2). Task-specific neuronal time constants can be learned, as demonstrated in [39, 40]; however, we here only assume a loose agreement between the timescale of the task and neuronal time constants, and thus set 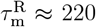 ms. Fig. 2 (right) shows that our model can solve the task using only six neurons (three representation, three error units).

### 3.2. Explicit error neurons scale to many areas, where dendritic error construction struggles

Our model differs from the bio-plausible models of error learning by Sacramento et al. [33] and dendritic hierarchical predictive coding (dendritic hPC, Mikulasch et al. [25]) through the existence of explicit error neurons, as in classical predictive coding models [28]. In Fig. 3, we demonstrate that our model scales to train many areas, and performs on par with an ANN. The networks are trained to learn an input-output mapping recorded from a randomly initialized teacher ANN with the same architecture; therefore, task complexity increases with the number of areas in the networks. This guarantees that the student networks are in principle able to solve the task, but also that good performance requires training of all areas – e.g., it is not sufficient to train only the output weights.

**Figure 3:**
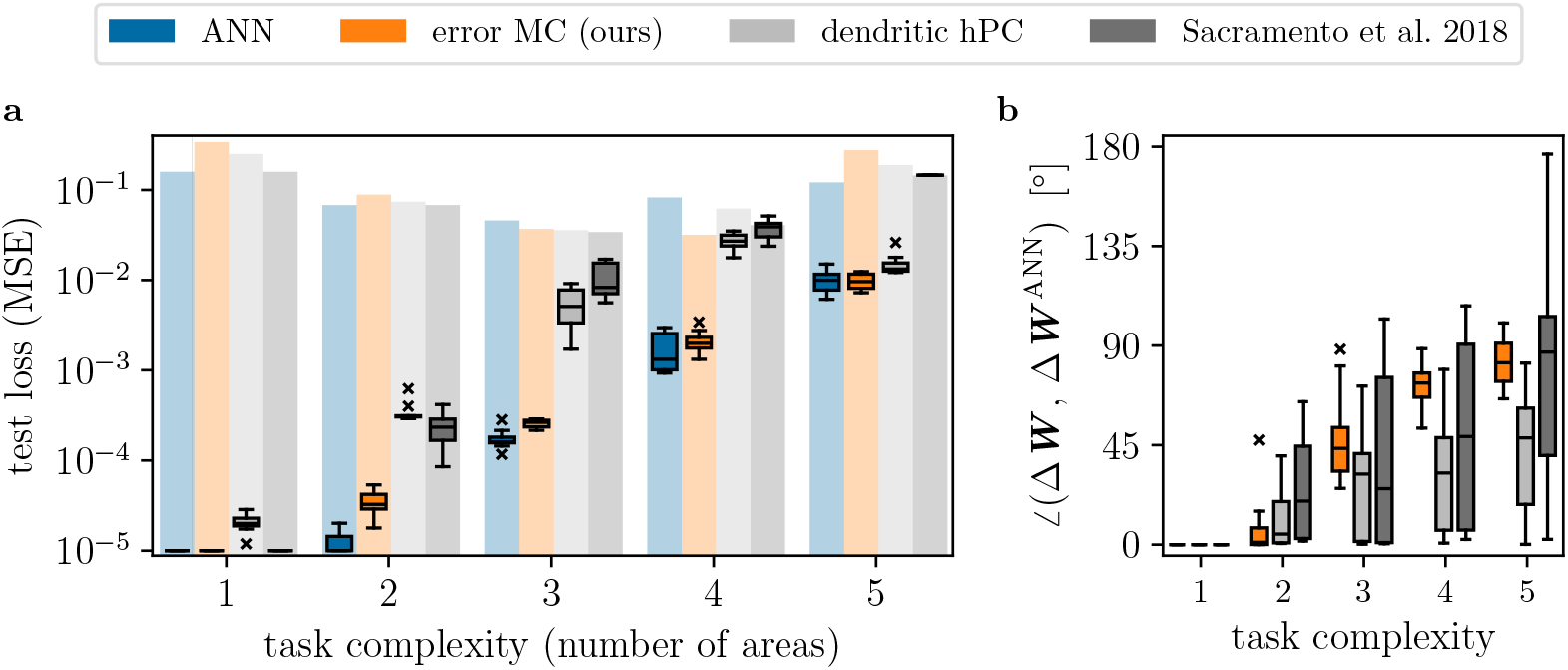
Error neuron microcircuit trains many areas, on par with an ANN. We compare learning in our model to an ANN trained with backpropagation, and the dendritic error computation models of Mikulasch et al. [25] and Sacramento et al. [33]. In order to show that our model scales to large networks, we systematically increase the number of areas. The task is to imitate a teacher network of the same architecture, where task complexity increases with number of areas. **a)** Mean loss on test set (median and quartiles over 10 seeds). Transparent bars in the background show test performance before training. Our error neuron microcircuit performs on par with the ANN, while the dendritic hierarchical PC model of Mikulasch et al. [25] and the microcircuit model of Sacramento et al. [33] decrease in performance when training more complex tasks. **b)** Alignment between synaptic weight changes in all three models vs. those of the ANN (median over all seeds and areas).

We compare performance to the dendritic microcircuit model of Sacramento et al.; due to the fundamental relationship between PCN and ANNs highlighted in Section 1, we are also able to include dendritic hPC in an inverted setup. To enable a fair comparison, we have extended dendritic hPC to a non-linear version in this work (see Appendix C); however, as we find that the non-linear version performs equal to the linear model (Fig. C1), we have used the original model of [25] here.

We find that the models with dendritic error construction do not scale well to multi-area tasks, indicating that dendritic error construction fails when errors are projected across more than two areas. On the other hand, our networks with explicit error representations always perform on par with the baseline, ANNs trained with backpropagation. This is an important result, indicating that current bio-plausible models of dendritic error computation lack the scalability required to explain learning across the cortical hierarchy (see Discussion).

Note that task complexity 5 cannot be fully solved by any network, as even the ANN baseline is only able to reduce the loss by about one order of magnitude. Indeed, training only the last three areas of the ANN achieves about the same performance, with a test loss of (1.2 *±* 0.1) *·* 10^−2^. This indicates that the networks are not able to make use of learning in all of their five areas, which explains why all models perform equally on this task (barring the model of Sacramento et al.).

Further, our model differs from the others by incorporating connections between all areas. To ensure that this is not the reason of its higher performance, we have repeated the experiment with strictly hierarchical connectivity only, see Fig. A1. Indeed, the networks with strictly hierarchical connectivity perform equally well.

### 3.3. Networks benefit from loose matching of error and representation units

An essential feature of our model are the populations of error and representation neurons in each area. Here, the precise one-to-one matching of prediction and error neurons usually found in PCN is replaced by preferential targeting, where local weights are not restricted to an identity matrix. As we will demonstrate, this does not only improve biological plausibility, but is also beneficial for task performance.

We train a two-area network representing the early visual areas (e.g. V1 and V2), with 784 representation and error neurons each in V1, and a variable number of *n*_*R*_ and *n*_*E*_ neurons in V2; see Fig. 4. Representation neurons are tasked to predict activity in V1 matching the label provided as top-down input. This generative task is therefore a direct implementation of predictive coding. The images fed into the network are 28 × 28 images of the MNIST dataset (one image per digit class).

**Figure 4:**
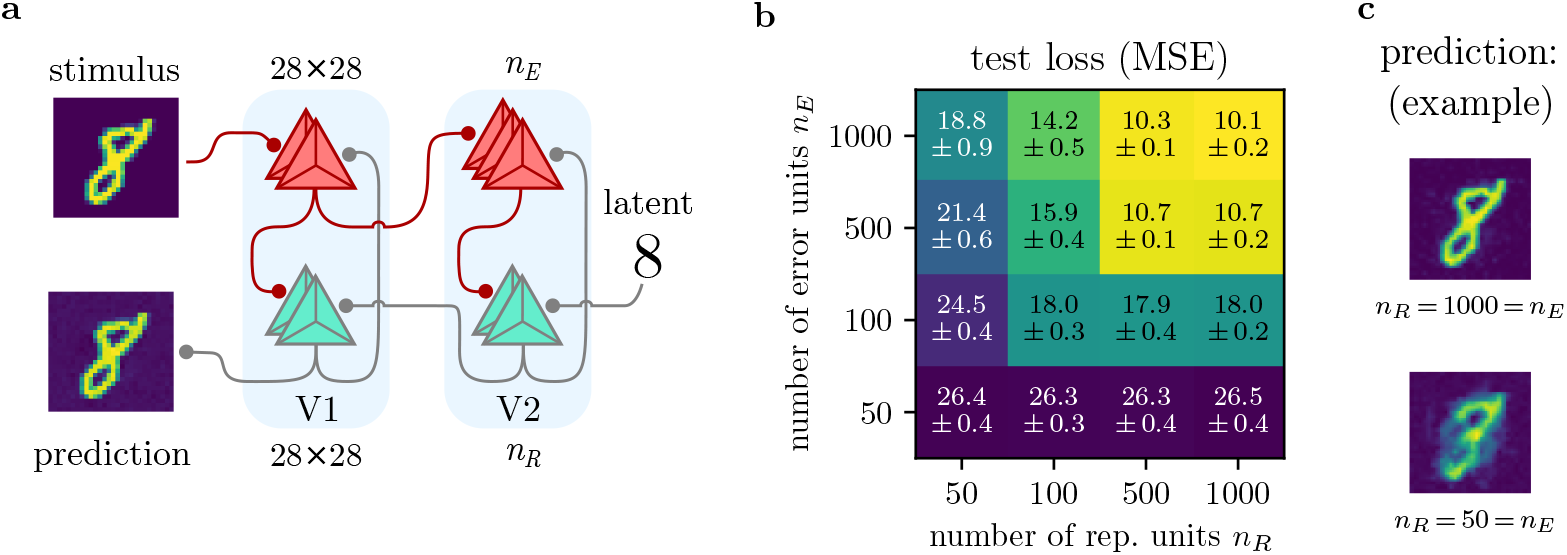
Implementation of predictive coding, and stability under variation of neuron populations. We demonstrate that our model accommodates predictive coding using a generative task, where the top-down input (‘latent’) is a digit [0,…, 9], and the desired prediction a 28 × 28 image from the MNIST dataset. **a)** Network setup. As opposed to previous tasks, the network is inverted, thereby implementing predictive coding. V1 is modeled with 784 representation and error neurons each, whereas in V2, the number of representation units *n*_*R*_ and error units *n*_*E*_ can be varied. **b)** Test loss for different *n*_*R*_, *n*_*E*_ (mean and stdev over 10 seeds). Performance generally increases with larger populations, but is stable even for *n*_*R*_ ≠ *n*_*E*_. Note that the networks make use of additional error units (*n*_*E*_ *> n*_*R*_). **c)** Example predictions of the digit ‘8’.

As Fig. 4 b & c show, the networks successfully learn to predict visual stimuli given a latent. To make sure that the network is actually required to propagate errors, we disable learning of *W* from the latent to V2, and indeed find drastically reduced performance with a test loss of 40.3 *±* 0.4 (*n*_*R*_ = 1000 = *n*_*E*_). Finally, an ANN trained on the task achieves similar performance to our model, for which the test loss is 9.2 *±* 0.1 (*n*_*R*_ = 1000).

As one expects, increasing the number of representation and error units in our model leads to better predictions (diagonal in Fig. 4 b). Interestingly however, the networks’ performance is also stable under variation of population numbers. For example, for all combinations of *n*_*R*_ = {500, 1000} and *n*_*E*_ = {500, 1000}, the mean square error (MSE) is approximately the same.

Notably, additional error units appear more useful than extra representation units: For a given number of representation units, increasing the pool of error units *always* improves performance, whereas more representation neurons are only beneficial if there already is a sufficient number of error units (*n*_*E*_ *≥ n*_*R*_). This implies an experimental prediction, where a larger pool of error units is generally favored over a larger number of representation units (see Discussion). Note that this is a feature enabled by loose matching of local populations: if each representation unit was assigned exactly one error neuron, additional error units could not project to representation neurons, and thus performance would not be improved for *n*_*E*_ *> n*_*R*_.

### 3.4. Error neuron microcircuits are robust to neuron variability and ablations, unless error signals become too noisy

To assess the impact of parameter noise and ablations on our model, we again consider the generative MNIST task of Fig. 4. First, we model variability of neuron activations *φ* in a network with [784-1000] representation units, and the same number of error neurons (Fig. 5 a; note that error units still are linear, 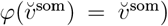. We introduce neuron-wise noise to the slope and offset/bias: activation slopes are noised by sampling from *𝒩* (1, *σ*_act_), where the standard slope used in the other experiments corresponds to 1. Similarly, offsets are noised using *𝒩* (0, *σ*_act_).

**Figure 5:**
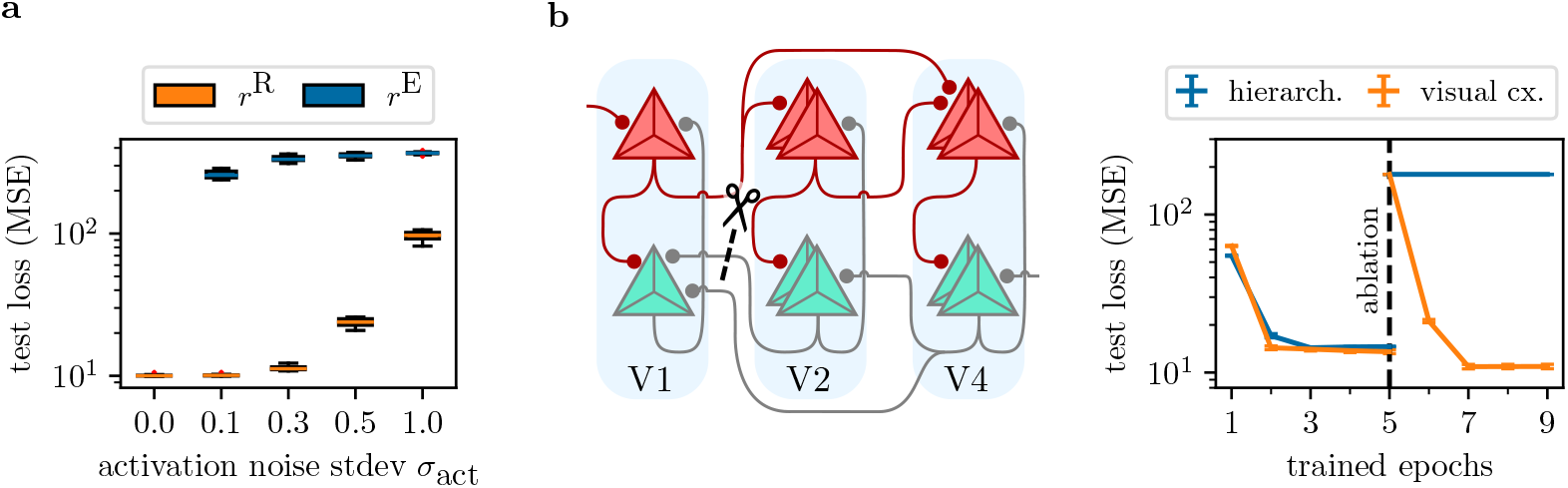
Effects of neuron variability and ablations under the generative MNIST task. **a)** We model neuron variability by adding parameter noise to the offset and slope of each neuron’s activation function. We sample from a normal distribution with standard deviation *σ*_act_ and mean 1 and 0 for slopes and offsets, respectively. Adding noise only to representation units (orange) shows their resilience even to large noise (e.g. *σ*_act_ = 0.3). However, as the networks rely on correct errors to be transported, they display high sensitivity to additional noise on error neuron activations (blue). **b)** We extend the setting of Fig. 4 to three cortical areas (V1, V2 and V4), and train two implementations of our model: one using strictly hierarchical connectivity, and one using the visual cortex connectivity with skip forward and backward connections (Fig. 1 b1). After initial training, we ablate connectivity from V2 to V1, and disable learning of new weights between these areas. While the hierarchical model is now unable to solve the task, the visual cortex model makes use of the skip connections to recover its performance. All points show mean and stdev over 10 seeds.

Fig. 5 a shows that network performance is rather stable under variability of representation neurons, *r*^R^ = *φ*(*ŭ*^som^). This can be explained by the algorithmic power of the error backpropagation algorithm: as long as errors are calculated correctly (no noise on 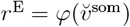), representation unit weights are trained to solve the task, independent of their activation function. Only for very large noise, the representation units are unable to learn the task in general, e.g. due to vanishing gradients caused by large slope factors (for which *φ*^*′*^(*ŭ*^som^) *≈* 0).

Noise on the error units on the other hand represents a fundamental challenge: In that case, representation units may possess the capabilities of solving the task, but are not guided to a solution due to faulty error signals. We see this reflected in the failure to train for any kind of noise on 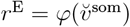. In Fig. A3, we demonstrate that this can be fully attributed to offset noise – the results can be interpreted using the framework of same-sign feedback alignment [69], for which we refer to the Discussion.

Finally, we study how ablations affect our microcircuit model (Fig. 5 b). In strictly hierarchical models of cortex, the disruption of synaptic weights between two areas fatally hinders prediction and learning. Due to our inclusion of realistic connectivity based on visual cortex, other areas may learn to take on the role of the ablated connectivity. We extend our model implementing the generative MNIST task to three areas, modeling V1, V2 and V4 with [784-500-500] representation neurons, and an equal number of error units. After initial training using the full connectome, we ablate the weights *W* projecting representations from V2 to V1, and disallow growth of new synapses between these areas. Upon continuing training, we see that the synapses projecting from V4 to V1 are able to take on the role of the ablated connections, and fully recover predictive performance. As expected, a strictly hierarchical implementation fails to do so, as there is no way for V2 or V4 to project to V1 anymore.

## 4. Discussion

### 4.1. Experimental evidence and predictions

Our model is based on recent experimental evidence and makes multiple predictions, the combination of which distinguishes it from other models of learning in cortex. It allows for two possible configurations: the (generative) predictive coding implementation, where a top-down stream aims to predict stimuli, and a classifier configuration with inverted streams (Fig. 1). Cortex may implement either configuration; however, due to the evidence of predictive coding in visual cortex, we will focus on the generative configuration in this discussion.

#### Explicit error and representation neurons

Like in the original model by Rao and Ballard [28], errors are encoded in mismatch neurons, and predictions are coded in a separate population of representation units. Following the canonical microcircuit for predictive coding, we hypothesize that both neuron types are pyramidal cells (PYR) found in the superficial and deep layers of cortex respectively [70, 71]. These populations are directly measurable, and separate our model from recent hypotheses like dendritic hPC [25] or the dendritic cortical microcircuits of Sacramento et al. [33], which assume populations of representation neurons inhibited by local interneurons.

Recent experiments have studied how errors are represented and used during learning, with particular focus on PYR in cortical layers 2/3 (L2/3), and their function of encoding prediction errors [12–19, 72]. In particular, motivated by evidence of local and inter-area connectivity (see below), we assume the error units to be feed-forward-projecting PYR in L2/3. Conversely, feedback-projecting PYR in L5 take on the role of representation units, and after learning or in absence of an instructive signal, the dendritic activity of these neurons is fully predictive of their somatic firing.

Based on these assumptions, our model predicts activity correlated with mismatch errors in superficial cortical layers, corresponding to the presence of error neuron populations, and to predictions coded in deeper layers, signaling the representation units of the model. While error units can be recognized by their decreasing activity during learning, representation units are not as easily identified: due to the recurrent structure of our model, representation units may code a mixture of prediction and mismatch. The degree of mixture can be varied by the conductance couplings {*g*^rep^, *g*^err^} of the two compartments (see Section 4.3). It may be the case that *g*^err^ is much larger than *g*^rep^; then, representation units mostly encode errors during learning, while their activity is fully explained by predictions in absence of mismatch signals. Indeed, this reflects the recent analysis of Aceituno et al. [73], showing that in-vivo activity of L5 PYR better aligns with strong influence of error signals (‘target learning’).

#### Error and representation neuron populations

Our simulations show that error neuron microcircuits can accommodate loose matching between the two populations within each area. In fact, we observe that it can be beneficial to recruit more error neurons to a task, whereas extra representation units does not change task performance (Fig. 4). It remains to be studied whether the ideal ratio of representation vs. error neurons given a fixed total number is 50%/50%, or it is always beneficial to have more error units.

#### Different dynamics of prediction and error units encode their roles

The two types of neurons in our model do not only encode different information, but also use different dynamics to perform their respective computations.

L5 PYR act as representation units as in the classical neuron model, summing and integrating all of their inputs in the soma [74]. Their two compartments can be realized by perisomatic integration zones, possibly as distinct basal and proximal apical dendrites. This allows for additive somatic integration, with only small sub-or supralinear effects (due to NMDA spikes).

The role of distal apical dendrites in representation units is not defined in our model. In [33, 75], it has been hypothesized that errors are coded in such distal apical integration zones. For error units in L2/3, this means that L5 PYR representation units may receive errors through apical dendrites reaching L1, in line with experimental data [19, 76]. Furthermore, experiments have shown that top-down or local L2/3 input to L5 PYR amplifies somatic activity [60, 77], e.g. by Ca^2+^-mediated plateau potentials. However, the mechanisms of such gain modulation do not allow for apical inhibition to reach the soma. It is thus unclear how L5 neurons can learn from negative errors, if they are coded exclusively in distal apical dendrites. In absence of such a mechanism, we hypothesize that distal apical dendrites encode additional information like top-down attention, uncertainty and awareness.

Error units, on the other hand, may make use of such a gain modulation mechanism to implement their multiplicative dynamics. Here, our model predicts that neuronal outputs of L2/3 PYR are formed by modulation of bottom-up errors with the activity of same-area L5 representation units: the concrete prediction of the model is that error neurons are silenced if activity of representation units saturates, *e ∝* (*r*^R^)^*′*^ = *φ*^*′*^(*ŭ*^som^). This could be realized by translaminar inhibition recruited by extreme values of L5 activity.

*Dendrite dynamics before/after learning*. In our model, the teaching signal from which representation units learn is encoded in an error compartment *u*^err^. Learning reduces this signal, until the representation compartment *u*^rep^ is fully predictive of somatic activity (see Fig. 6, left). As shown in Fig. 1, the two compartments may be realized by separate apical/basal or perisomatic dendrites.

**Figure 6:**
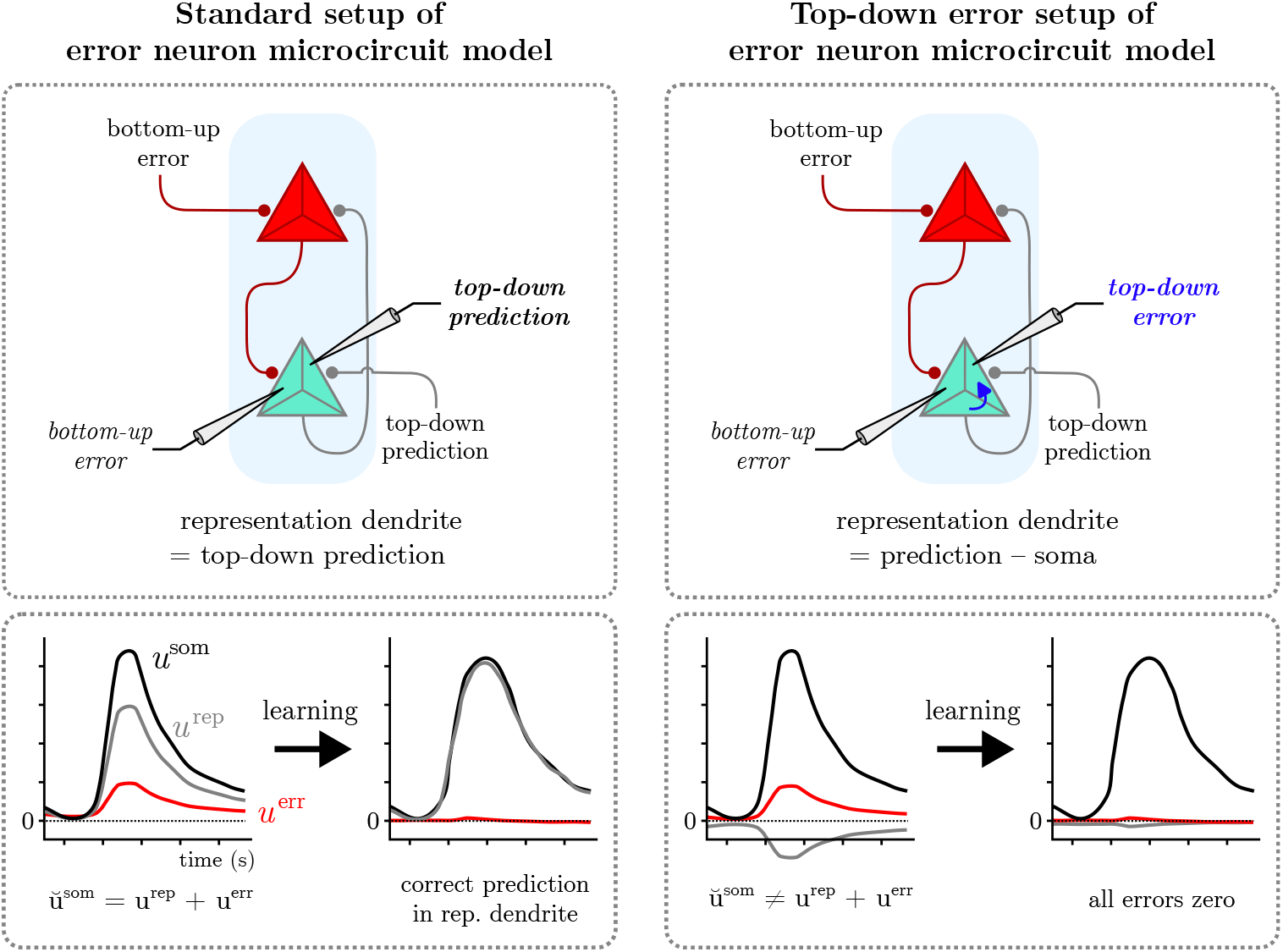
Alternative description of our model with top-down and bottom-up errors in representation and error dendrites, respectively. Classical predictive coding models describe two errors in each area (top-down and bottom-up). In the standard formulation of our model (left), each representation neuron encodes only bottom-up errors (in its error dendrite); however, the model admits an alternative setup, where predictions are compared to somatic activity on the representation dendrite (right). These two interpretations are computationally fully equivalent, with equal somatic and firing activity as well as learning performance (see Appendix D). The only difference lies in different measurable correlates: in the standard setup, activity in representation dendrites is never zero in the presence of top-down predictions, while the soma integrates top-down predictions with bottom-up errors. In the top-down error setup however, representation dendrites ‘explain away’ the difference between soma and top-down predictions, leaving only a residual top-down error, which is minimizes during learning. We stress that *u*^som^ remains the same in both interpretations.

A recent study of Francioni et al. [75] has analyzed this scenario using optogenetic measurements of L5 PYR in mice performing a brain-computer interface (BCI) task. The authors assign two randomly chosen populations P^+^/P^−^, depending on their contribution to solving the task (rotating a bar from 0 to 90 degrees by matching a neuronal target activity). Indeed, they find that the difference between somatic and distal (!) apical activity correlates with mismatch signals, where the two populations integrate positive and negative error respectively. The methods applied in this work are highly innovative and pave the way for quantitative studies of error learning on the neuronal level, however several questions remain to be clarified. In the study, the target is defined as a specific difference in activity P^+^− P^−^. Here, the BCI setup may lead to a confounding of learning and co-activated neurons, as the P^+^/P^−^ populations may include task-unrelated neurons. Furthermore, the task is highly stereotypical, as angle initialization and target are always defined as exactly 0 and 90 degrees respectively, and the target is only approachable from one direction (target > angle always, i.e. task is asymmetric). Thus, the setup is susceptible to habituation, and does not probe all possible error signals (target > angle vs. target < angle); under the ‘vector hypothesis’ (i.e., backpropagation), one expects to see a sign flip of the error for both populations when changing the direction of approach. It remains to be studied whether their findings extend to more general task setups.

One *apparent* distinction of our model is that there is only one type of error, the one encoded in error unit somata and communicated locally to compartments *u*^err^ of representation units. Many implementations of PCN compute two errors in each area: bottom-up and top-down errors [25, 28, 65, 78]. For example, in [25] (Box 2), basal errors *b* encode the usual learning signals (exactly as our *u*^err^), while apical errors *a* are formed by the difference between afferent representations and somatic activity (no direct correlate in our model). This implies different experimental signatures, where basal and apical dendrites both code error signals, which should decrease their activity during learning.

However, note that after convergence of somatic dynamics, both errors are equal up to their sign, *a* = *−b*, in dendritic hPC (cf. Eq. XI in [25]). I.e., after transient dynamics, only one error signal is in fact encoded, the same in both compartments. Our model allows for an alternative formulation, where top-down errors are also computed on the representation dendrite (see Fig. 6 right). As we demonstrate in Appendix D, our model can be rewritten in order to reflect this exactly, and is thus computationally equivalent to the standard formulation, while retaining its scaling advantage over dendritic hPC (cf. Fig. 3).

Based on this reformulation, our model (and dendritic hPC) allows for two differing experimental predictions: after transient dynamics have settled, either representation and error compartments encode predictions and errors respectively, or they both code the same error signal (with opposing signs, and up to rescaling by conductance couplings), which decreases during learning. Note that the somatic activity is the same in both models in either case (soma = sum of all afferent inputs). An issue of the alternative formulation is the need for the somatic potential to act *subtractively* on the error dendrite compartment (blue arrow in Fig. 6) – it not clear how biology can implement this, which is why we have chosen to model only one type of error compartment per representation unit in the standard formulation of our dynamics.

#### Inter-area connectivity

Our model differs from classical PCN (e.g., the Rao-Ballard model [28]) by its inter-area connectivity scheme, see Fig. 7. Our configuration has multiple benefits in terms of computational power and biological plausibility of the neuronal calculations, as we describe in Section 4.2.

**Figure 7:**
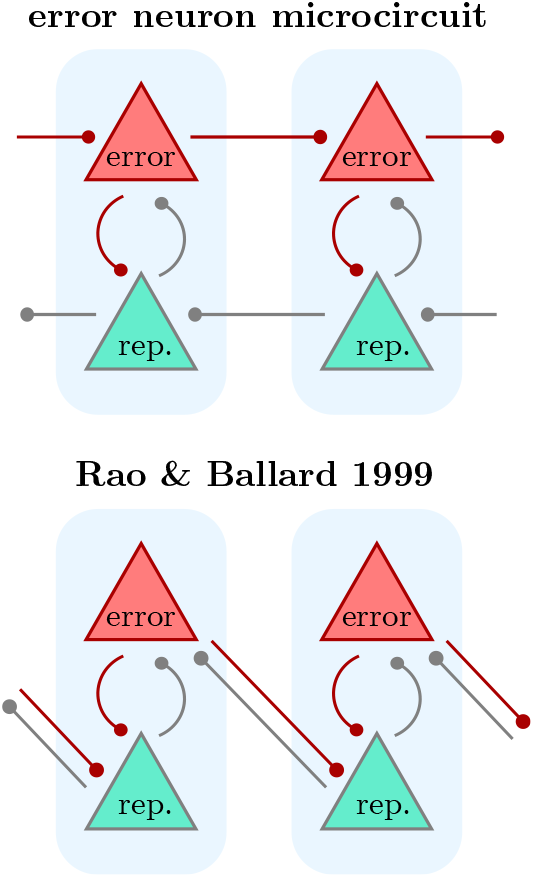
Connectivity in our model and classical PCN.

In detail, our model predicts two separated streams of errors and predictions. Various studies have investigated preferential targeting of dendritic compartments by top-down/bottom-up projections [15, 16, 18, 48, 74] and mapped out distinct feed-forward and feedback pathways in the cortical hierarchy [53].

Concretely, if we assume that representation units are PYR in L5, and errors are encoded in L2/3, this predicts a top-down stream of L5 *→* L5 projections, and an opposing error stream projecting L2/3 *→* L2/3.

However, it appears unlikely that cortex implements the *exact* connectivity of our circuit, or for that matter of the Rao-Ballard model. Instead, we have to interpret the inter-area connectivity as effective functional connections. For example, a recent analysis by Balwani et al. [72] of visual behavior in mice shows functional connectivity accommodating our circuit and the Rao-Ballard model to varying degrees (see Fig. 5A therein): novel images induce strong feed-forward projections from L2/3 *→* L4 (V1 to LM), which may be passed locally to L2/3 of LM, in agreement with strong same-area projections from L4 to L2/3 in the canonical cortical microcircuit [70, 79–82]. Such effective functional connectivity may represent our L2/3 *→* L2/3 error stream.

In the analysis of [72], the feedback stream (LM to V1) of L5 neurons shows projections both to L5 but also L2/3; in this direction, it appears that neither our and nor the Rao-Ballard connectivity are cleanly observed, but rather a mixture of both. In our model, this may be explained by L5 representation units mixing predictions and errors (non-weak nudging, ‘target-learning’), confounding the distinction of functional projections of error and representation pathways. Alternatively, our model can be modified such that representation units do not project locally to error units in the same area, but to error units in upstream areas.

The exact nature of functional connectivity during inference and learning remains to be studied; however, the clear roles of the *effective* L5 *→* L5 top-down and L2/3 *→* L2/3 bottom-up streams are testable predictions of our model.

#### Local connectivity

As in the Rao-Ballard model, our model assumes same-area connectivity between error and prediction units, which may be implemented by L2/3 and L5 PYR. Local connections between these populations are well characterized, see [83] for a recent review.

L2/3 neurons receive strong projections from L4, which we interpret as the inter-area pathway of errors (see above). However, L2/3 neurons (in particular L2) additionally receive strong input from PYR of L5A [84–87], in line the input to *v*^rep^ with our model. Additionally, L4 neurons receive numerous inputs from local L6 PYR (see e.g., [88]), whose role has been proposed to be modulatory [89, 90]. Experimentally, our model predicts that L2/3 error neurons receive bottom-up errors which are gain modulated by local L5 inputs, and that saturating representation units silence error neurons. We hypothesize that this gain modulation may be performed by plateau potentials, analogously to L5 PYR [60], or non-linear dendritic dynamics of the L2/3 neurons.

Computationally more important however is the local connectivity of L2/3 *→* L5, as this communicates the error signal to representation units. Strong connectivity between these areas has been observed [80, 83], and it is part of the canonical cortical microcircuit [79, 91].

Our model predicts that these projections are crucial for learning. For example, if the relevant L2/3 *→* L5 projections are inhibited, then L5 PYR only integrate top-down activity, and there is no differential local error signal from which these units can learn.

For all local connections, we have assumed loose matching between error and representation units. For simplicity, we have modeled preferential targeting as one-to-one matching (identity weight matrix) with noise, effectively connecting all representation and error neurons more or less strongly. More concretely, biology may implement an exponential distance rule of local axonal arbors, i.e. local connectivity *L*^RE^ and *L*^ER^ scaling as 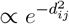, where *d*_*ij*_ is the distance between neurons *i* and *j*. Such scaling has been observed in inter-area conenctivity in different species [95–97], and potentially also applies to local connectivity.

#### Noise sensitivity

The results of Fig. 5 a show that error neuron microcircuits are largely insensitive to variability of representation units, but noise on errors easily disrupts learning. We have identified that this is due to noise on the offset of error units (Fig. A3), changing the sign of error signals. Algorithmically, this makes sense: it is known that successful learning requires errors of the correct sign, whereas the magnitude is not as important (see same-sign FA [69, 98]). Therefore, our model makes the experimental prediction that perturbing error units disrupts learning much more than perturbing of representation units, which can be tested by injecting the same noise levels to L2/3 and L5 neurons respectively.

### 4.2. Embedding into cortical models of learning and predictive processing

Most implementations of error learning in the cerebral cortex can be grouped into two categories: those with error neurons (as in the classical Rao-Ballard model [28]), and models where errors are constructed locally on the dendrites of representation units. In the implementations of dendritic error computation by Mikulasch et al. [25] and Sacramento et al. [33], errors are constructed as the difference between prediction and lateral inhibition on the representation units’ error dendrites; for a summary of the principle, see Fig. 1 in [25]. This implies that representation neurons compete through lateral inhibition, adding biological credibility to such models, and send their predictions both up and down the hierarchy, without the need for two opposing error/representation streams.

However, dendritic error construction is not well tested as a model of the cortical hierarchy, and scaling to multi-area networks remains to be proven. In [33], Sacramento et al. successfully trained a three-area networks on the MNIST digit classification task; however, the same performance can be reached by leaving out the first area [99], making it unclear if useful error signals have reached the early area in these results.

We claim that explicit error neurons facilitate more stable learning across cortical areas, as dendritic error construction is susceptible to noise, caused by imperfect lateral inhibition. In short: explicit error neurons allow errors to be propagated through the network *and then* sent to representation units, while dendritic error construction requires the calculation of a difference of activity *in every area*. It is precisely this difference which is propagated as the error; under noisy weights and neuron dynamics, this reconstruction of useful error signal fails, as Fig. 3 shows.

Consequently, the most important computational achievement of our model is that it scales to many areas, while maintaining all of its bio-plausible constraints. Many studies in the literature have significantly simplified their model dynamics when scaling up simulations, e.g. by assuming ideal weight sharing [25, 33], replacing neuron dynamics by steady-state approximations or artificially re-initializing voltages [32, 33, 100], or implicitly removing recurrency by calculating separate forward and backward passes [33, 101]. Here, we have demonstrated that our model is able to scale without changing the implementation of its dynamics in any way.

Due to the equivalence of predictive coding and backpropagation under the ‘fixed prediction’ assumption (see Section 4.3), our model is also an implementation of predictive processing. In particular, for ideal local weights, it is computationally equivalent to classical PC models [28, 65], but differs in its inter-area connectivity and neuronal dynamics. Most apparent is that we have two separate prediction and error streams, while classical PCN have bidirectional projections between error and representation units (Fig. 7). Why do we implement such connectivity? Our model has a number of advantages:

i. Faster and less noisy: predictions don’t need to go through error units in our model, which would double the number of neurons involved in an inference step – this affects processing speed and adds noise sources.
ii. Simple switching between inference and learning: as the error and prediction streams are only loosely coupled, error signals can be introduced dynamically, without the need to wait for the dynamics of all neurons to re-settle. This may be particularly important for tasks where learning signals are only provided at discrete times, such as reinforcement learning.
iii. Signal transport and local weights: in most PCN, error units compute the local error as a difference of voltages, 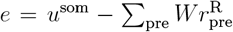, where *u*^som^ is the somatic voltage of the prediction unit in the same area (Fig. 7) [21]. Without additional mechanisms, it is unclear how error units obtain *u*^som^ in the setting of non-linear neurons and non-identity local weights. To solve the former issue, errors can be defined on the rate level, such that 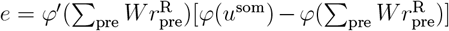 as in the Rao-Ballard model [28]; but then it is not clear how error neurons can calculate *φ* and *φ*^*′*^ using *exactly* the activation function as local representation units. In either case, it appears that error units serve as pseudo-compartments of representation units instead of actual neurons. In our model, errors are explicitly projected from area to area by the error neurons, without the need for local error computation. These errors are then projected locally to representation units, which make them available at the synaptic site *W*.

### 4.3. How closely do we need to implement ‘backpropagation and the brain’ ?

BP is the algorithm underlying the deep learning revolution, and in practice currently the only way ANNs are trained to solve complex tasks. While alternatives such as reservoir computing have been proposed [102, 103], they fall short in terms of scalability, complexity of learnable tasks, and efficiency. For example, reservoir computing requires up to 60–120 times as many neurons to reach the same predictive performance as an LSTM network trained with backpropagation-through-time (BPTT) [104], and feed-forward networks with frozen hidden weights perform significantly worse given the same number of neurons [105]. Assuming that the cortex has evolved to use fewer neurons and thus consume less energy to achieve more, this is what motivates the ‘backpropagation and the brain’ hypothesis.

While it is clear cortex does not implement the error backpropagation algorithm *exactly* as it is formulated in machine learning, one may explore the hypothesis that some approximate version can explain learning [1, 2]. In our model, backpropagation is demonstrably obtained in the limit *g*^err^ *→* 0, where error and prediction pathways decouple, in conjunction with strictly hierarchical connectivity and identity local weights (Section 6.2). While the inclusion of such deviations from an ‘ideal’ implementation of BP is motivated by biological plausibility, it is important that such properties can also be interpreted as features, e.g. as implementations of advanced machine learning techniques. Thus, we now discuss how both non-hierarchical connectivity and non-weak nudging 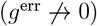 can fit together with backpropagation on a firm theoretical understanding.

Loose hierarchy and recurrency: Following connectivity in the Macaque visual cortex, we have organized connections in our model into preferential hierarchical streams, but also include skip connections. More generally, other modalities may implement less hierarchical organization, as non-primate species do in general [97, 106–108]. At first glance, this seems at odds with the success of hierarchically organized ANNs, and one may assume that this excludes any comparison between feed-forward ANNs and biological networks. Algorithmically however, skip connections can be made sense of through their relation to residual networks [109, 110], which draw their computational power from the central motif of identity (skip) projections.

Note however that the vanilla BP algorithm is derived for strictly hierarchical networks only (layered graphs), where the activity of each neuron is causally dependent only on input from earlier areas, and can be calculated in a closed form. This can be extended to include skip connections (directed acyclic graphs), as we do in this work. However, for general, recurrent architectures including loops and self-recurrency, one needs to take the dependence of each neuron on past network states into account. To do so, one can generalize BP to the backpropagation-through-time (BPTT) algorithm. This solves the issue by unrolling the recurrent network in time to obtain a layered graph, increasing however biological implausibility. Online learning approximations to BPTT have been proposed [40, 111–113], and future work may investigate microcircuit adaptations of these algorithms. In particular, the solution in [40] incorporates the same representation/error neuron populations presented in this work, thus lending itself to a cortical microcircuit implementation.

Non-weak nudging: While classical implementations of PC require representation units in the first area to be clamped to the stimulus, our model allows for a more organic introduction of targets into the network. Instead of clamping, it couples error and representation units with a conductance parameter *g*^err^, with the ability to implement both weak nudging (*g*^err^ *→* 0) [41] and strong nudging, restoring clamping in the limit of very large *g*^err^. As a result, representation units across areas encode a mixture of bottom-up errors and top-down representations. This has important implications for experimental predictions: In a strict interpretation of ‘backpropagation and the brain’, one expects to find that representation units do not encode errors at all (‘fixed prediction’ assumption [23]). This begs the question how representation units can access error information in order to update their weights, and is at odds with recent experimental evidence by Aceituno et al. [73]. In fact, the results of [73] suggest that representation units are strongly influenced by targets during learning, aligning with the clamping hypothesis. In our setting, varying the conductance *g*^err^ provides an organic way to shift from backprop-like (weak nudging [32, 41]) to target-like learning (clamping).

Coincidentally, theoretical work on bio-plausible backpropagation has demonstrated successful training on larger datasets even for strong nudging [100, 114, 115]. Intuitively, by changing the influence of errors on the network, representation units change the relative mixture of how strongly they encode bottom-up and top-down information, and weights may follow a completely different trajectory than that of strict backpropagation; we observe this effect in our model in Fig. A1 b, where networks train well even as their weight updates differ up to 90^*°*^ from backpropagation.

In conclusion, this demonstrates how advanced machine learning techniques can inform models of learning and connectivity, and shows that ‘backpropagation in the brain’ implies a landscape of theories with distinct degrees of bio-plausibility.

### 4.4. Future work

Several issues remain unaddressed in this work, some of which we have highlighted in the previous sections. Additionally, the following points are left for future work:

- Unsupervised and reinforcement learning: we have tested our model on supervised benchmarks, which require external target labels. In the framework of predictive processing, this means that we have provided a fixed, sharp prior of expectation for every stimulus. However, by replacing the loss function, our model can also accommodate other learning schemes, where latent representations are formed purely by observation, or only sparse rewards are provided. In particular, it could serve as a microcircuit implementation of theories of decision-making, action selection, and motor control, such as active inference [116] or reinforcement learning in the cortico-basal ganglia-thalamic loop [117–120].
- Spikes: to extend our model to spikes, spiking AGREL [45] and a recent spiking implementation [121] of the dendritic error construction model [33] can serve as guidelines.
- Signed errors: in our model, error units have linear activations, i.e. they can produce both excitatory and inhibitory outputs. This issue has been discussed in the literature, with a range of possible solutions based on separate populations for negative/positive errors, error signals coded with respect to a bias, or transformed errors [61–65]; see [122] for a comprehensive summary.
- Dale’s law: neurons in our model can change their weights from excitatory to inhibitory, and vice versa, in contradiction with biology. Recent works have successfully trained networks of purely excitatory and inhibitory populations [72, 123], but rely on standard training libraries without a mechanistic explanation of the error propagation. It remains an open question how to model error neuron microcircuits observing Dale’s law.
- Connectivity: in our description, we are singling out the interconnectivity of feedback-projecting PYR in L5 and feed-forward-projecting L2/3, excluding relevant projections from thalamus and L3B as well as L6. While this is motivated by the high degree of connectivity between L5 and L2/3, our model will need to be embedded into a framework accounting for all relevant projections – in particular higher-order thalamic inputs, given their strong influence on L5A and L1 [124], and their computational relevance [125–130].
- Recurrent populations: while our model includes recurrent dynamics, local populations of error and representation units are not self-connected. Conversely, real PYR in L5A are highly interconnected [80, 131], prompting a generalization of our model to recurrent populations.
- Active dendrites & temporal dynamics: while our model makes use of slow membranes to encode short-term memory, and non-linear dendrites to estimate the gain factor of errors, it doesn’t account for the full complexity of neuronal dynamics. Future work may investigate fast top-down prediction and slower error signaling, which produce distinct time scales in experimental signatures [72].

## 5. Conclusion

The ‘backpropagation and the brain’ hypothesis remains controversial, but as we have argued, the discovery of error units in cortex and the assumption that the brain uses existing errors to their fullest makes it a compelling theory to study. An important milestone in this regard is the realization that predictive coding can be formulated as an approximation of backpropagation along arbitrary computational graphs [21–24].

Our model aims at filling the gap between deep learning and neuroscience: it harnesses the proven functionality of hierarchical ANNs (where deep learning is established as the state of the art), and extracts connectivity and microcircuit motifs which guide a bio-plausible model. This top-down approach separates our model from bottom-up approaches (e.g. E/I balance networks or Gabor filter models), which are difficult to functionalize [1, 132] or rely on unexplained, external learning mechanisms such as the surrogate gradient method.

In this work, we have shown that the error neuron microcircuit model can closely implement the backpropagation algorithm, and that such networks perform well and scale to solve complex tasks requiring learning across multiple areas. We hope that future work will clarify whether, and to which degree, the brain implements a form of error backpropagation.

## 6. Methods

### 6.1. Bio-plausibility of the error backpropagation algorithm

In a hierarchical ANN with *𝓁* = 1,…, *N* layers, representations are projected through weights *W*_*𝓁,𝓁−*1_, and layer-wise errors can be obtained recursively from a target provided to the output layer *N*. The error backpropagation algorithm, applied to such an ANN learning a non-temporal task, consists of two computational steps: the backpropagation of errors,

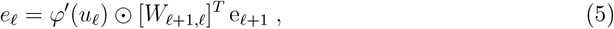

and the weight update rule

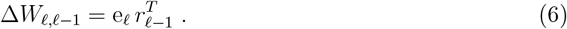

Implementing these computations into a bio-plausible model carries several obstacles [1, 2]. Our model solves the most pertinent ones in the following way:

**Table 1:**
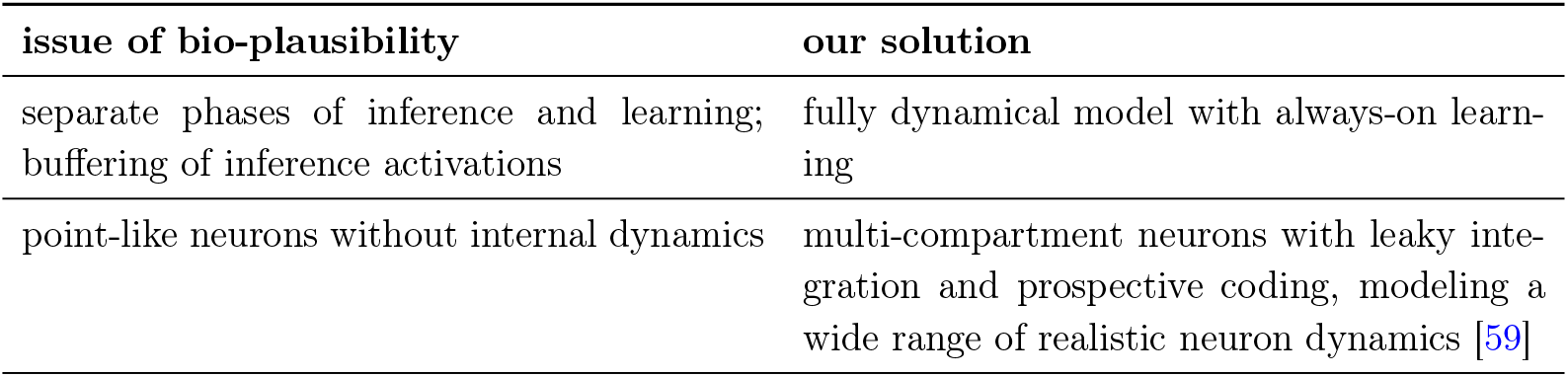
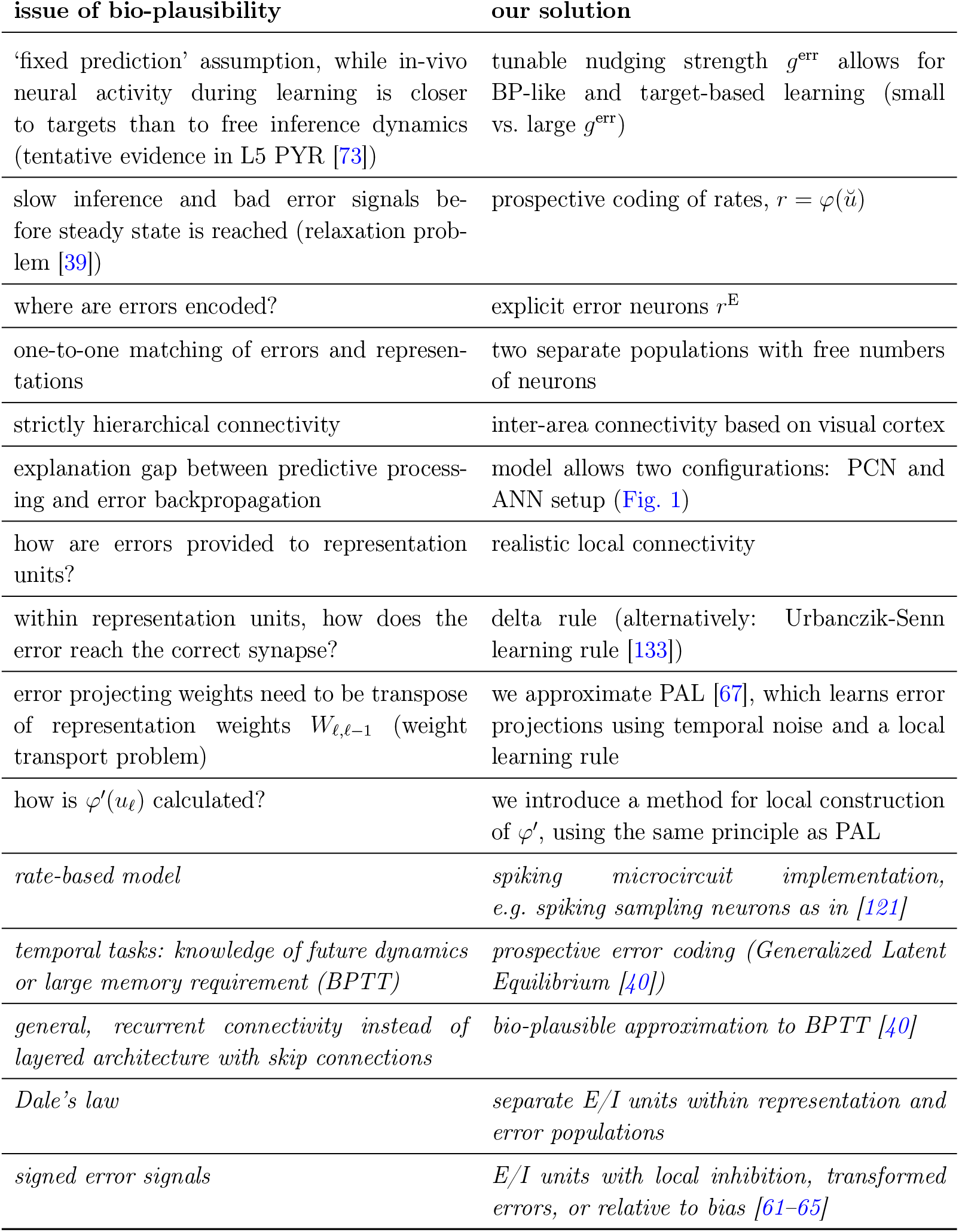
Translation of error backpropagation to bio-plausible model. The *last five entries* show important issues not addressed in this work, and potential solutions.

### 6.2. Neuronal dynamics and learning rule

Our microcircuit model implements Equations (5) and (6) in the following way. We define a network with *𝓁* = 1,…, *N* areas, equivalent to layers in the context of ANNs. In our model, we relax the strict hierarchical connectivity, and allow for connections between all areas.

The soma dynamics of the population of representation units in area *𝓁*, written as a vector 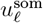, are

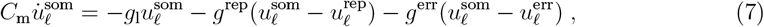

that is, a leaky integration of the compartment voltages 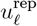and 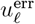. Compartments ar modeled as

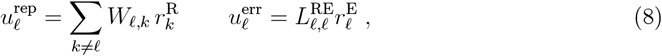

where *W*_*𝓁,k*_ represents the matrix of weights connecting area *k* to *𝓁*. 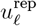 sums all inputs from representation units in other areas (but not from the same area), whereas 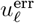 is formed from output of error neurons in the same area.

The neuronal output is the ‘prospective’ rate [39]

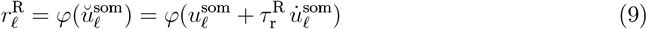

with 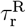 a neuron-wise parameter, called the prospective time constant. For quasi-instantaneous transmission, we set 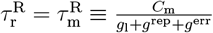. For tasks requiring memory, we encode short-term memory through slow neuron responses, for which 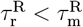 (see Section 6.5 and Appendix B).

Error units follow the dynamics

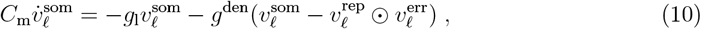

With

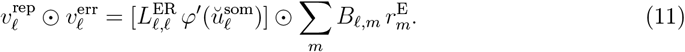

Note that 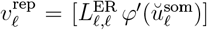 does not carry the units of a voltage, but rather 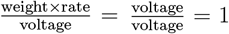, i.e. it is unitless. Thus, the summand 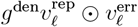 in Equation (10) is a current, and can be added to the somatic integration.

For learning, we need to inject a target signal into the network. We do so in the form of a target rate *r*^tgt^, which is provided to area *N* (‘output layer’) following slightly modified dynamics:

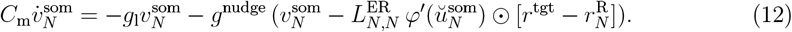

This means that the representation neurons in area *N* communicate their rate 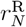 locally to error units, which calculate the difference between *r*^tgt^ and 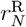.

Error neurons also implement prospective rates,

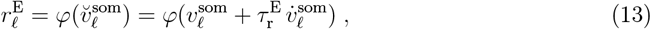

and we set all error neurons to be instantaneous, with 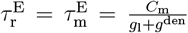 for hidden areas; 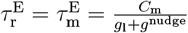 for the output area. Note that in all of our simulations, we choose a linear activation for error units, 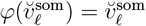.

By plugging Equation (11) into Equation (10), we can see that 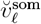 encodes the error in each area:

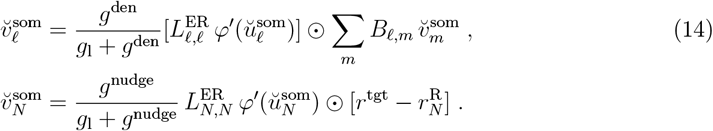

In a hierarchical setup with ideal local weights, 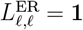, this implements Equation (5), under the assumption that 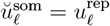 (see below).

The dynamics of the representation-projecting weights *W*_*𝓁,k*_ are given by

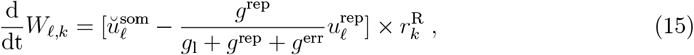

where by comparison with Equation (4) we can identify the total conductance as *g*^tot^ *≡ g* + *g*^rep^ + *g*^err^. By rewriting 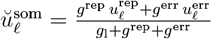, we obtain

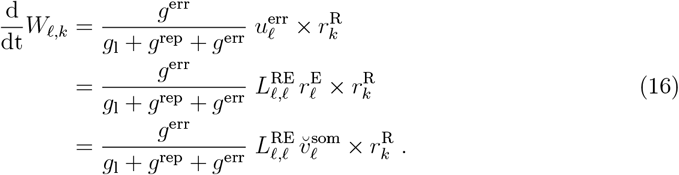

For identity local weights and hierarchical connectivity, we find that this implements Equation (6).

However, there is one additional caveat to the exact correspondence of our model and backpropagation: due to the coupled compartments, the soma of representation units does not only encode external inputs, 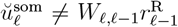. Instead, errors are added to somata in every area, confounding the activity of all units as a combination of inputs and errors due to the recurrency in the network. BP is only strictly equivalent to our model and predictive coding implementations under the ‘fixed prediction’ assumption, i.e. where the feed-forward activity 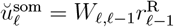 needs to be saved to be used during learning.

This is reflected in the angle of alignment between Δ*W* in ANNs and our model

(cf. Fig. 3 b), which tends to be larger than zero. This can be ‘remedied’ by decreasing the influence of the error compartment on the soma, *g*^err^ *→* 0, a limit also known as weak nudging [41]. However, this also shrinks the learning signal, as evident from Equation (16), which in turn leads to higher noise sensitivity and is less faithful to tentative in-vivo evidence [73]. Instead, we keep *g*^err^ as a free parameter; as our results show, our model performs on par with an ANN even for large nudging parameters, implying that neuron activities do not need to exactly follow those of feed-forward ANNs for successful learning.

### 6.3. Network connectivity

#### Visual cortex connectivity

To accurately model the degree of hierarchy found in cortex, we base the inter-area connectivity of our network on the projection densities found among the first areas of the Macaque visual cortex. We use the weighted connectivity matrix determined by Markov et al. 2014 [46] (Fig. 11 b therein, first six areas reproduced in Fig. 1 b1), measured by retrograde tracing.

To model the learning of a new task based on an existing network structure, we initialize each network using the relative connectivity strength found in Fig. 1 b1. For example, weights between V2 *→* V4 are larger than those between V2 *→* TEO by a factor of about 3 at initialization. We round all projections below 10^−2^ down to zero, and set the diagonal of Fig. 1 b1 to zero, as the recurrency within an area is already encoded in the local connectivity motif (local communication between representation and error units).

#### Weight symmetry

In our networks, errors are projected through weights *B*_*𝓁,m*_. Exact error backpropagation requires symmetric weights between two areas, i.e. *B*_*𝓁,m*_ = [*W*_*m,𝓁*_]^*T*^. While networks can learn even with non-symmetric, static feedback weights (Feedback Alignment, FA [66]), this has been shown to not scale well to complex problems, requiring more neurons to successfully learn a task [29, 67, 98, 134, 135].

An efficient mechanism to learn symmetric weights between two areas is Phaseless Alignment Learning (PAL) [67]. PAL uses temporal noise as an information carrier, allowing to simultaneously learn all error projection weights *B*_*𝓁,m*_, without disruption of representation unit weights *W*_*m,𝓁*_. Note that PAL makes use of the same principles as the mechanism for local reconstruction of *φ*^*′*^ proposed in this work, which we explain in Section 6.4.

In [67], we have shown that PAL is able to efficiently train the dendritic microcircuit model of Sacramento et al. [33]; however, its simulation on conventional hardware is time-consuming due to the requirement of fine-grained simulations of temporal noise.

For computational feasibility, we therefore approximate PAL by setting error projecting weights to the transpose of representation unit weights, and adding noise to mimic the approximate degree of alignment typically found in networks using PAL. We also need to consider that the numbers of error and representation units are not required to be equal in our network. To account for this, we pad the weight matrix with zeros: for example, 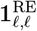 is an *n*_*R*_ ×*n*_*E*_ matrix (*n*_*R*_: number of representation units in area *𝓁*; *n*_*E*_: number of error units), with the ‘diagonal’ entries set to 1, and all others set to 0. For example, for *n*_*R*_ = 2 and *n*_*E*_ = 3 in area *𝓁*, we have

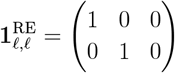

After every time step, error projections are set to 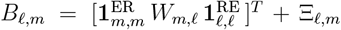, with Ξ_*𝓁,m*_ ∼ 𝒰 (−0.5, 0.5) sampled *once* (i.e. at the initialization of the network).

#### Local connectivity

In our model, we relax the strict one-to-one matching of error and representation units found in many models [24, 28, 33, 39, 136]. To model noisy preferential targeting between representation and error units, we fix local weights to 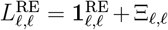, with Ξ_*𝓁,𝓁*_ *∼ 𝒰* (*−σ*_*L*_, *σ*_*L*_), and similarly for *L*^ER^. See Section 6.5 and Appendix B for all simulation parameters.

### 6.4. Non-linear dendrites enable local computation of φ^′^(u) in error units

Here, we explain how the error neurons reconstruct 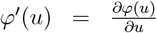 in their dendritic compartment. To simplify notation, we use a generic voltage *u* in this subsection, acting as a stand-in for the prospective voltage *ŭ*^som^ of the representation units.

Error neurons are able to estimate *φ*^*′*^(*u*) using biologically plausible ingredients: temporal noise, which is ubiquitous in the brain, and simple functions (high-pass filtering, gating and temporal averaging) that a dendrite can perform.

To start, we take a local population of representation neurons with membrane potential vector *u*, and add noise fluctuations for each neuron, *u* + *ξ*, where *ξ* can be e.g. white noise; note that any noise with sufficiently short correlation time suffices. In order for the neurons to encode a useful representation, the signal-to-noise ratio needs to sufficiently high, meaning that |*u*| *≫* |*ξ*|. Coincidentally, this is a useful property which we exploit in the following: The error neuron receives the noisy output of the representation unit at its dendrite,

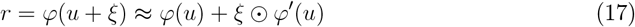

where we have expanded the rate to first order in small noise *ξ*. Next, we aim to isolate the component of *r* governed by noise (second term). Note that *u* is driven by stimuli and error signals in the network, which are of low frequency compared to the fast noise *ξ*. Therefore, we hypothesize that the error neuron dendrites integrate a high-pass filter, leading to

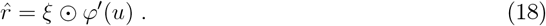

By applying non-linear gating onto its input, e.g. in the form of a rectified linear (relu) function, and averaging over noise samples, the dendrites compute

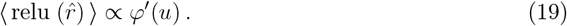

This shows that a rescaled version of *φ*^*′*^(*u*) can be computed locally in the error neuron dendrite, for sufficiently smooth activation functions. See Fig. 8 for simulation results for one pair of representation and error units.

**Figure 8:**
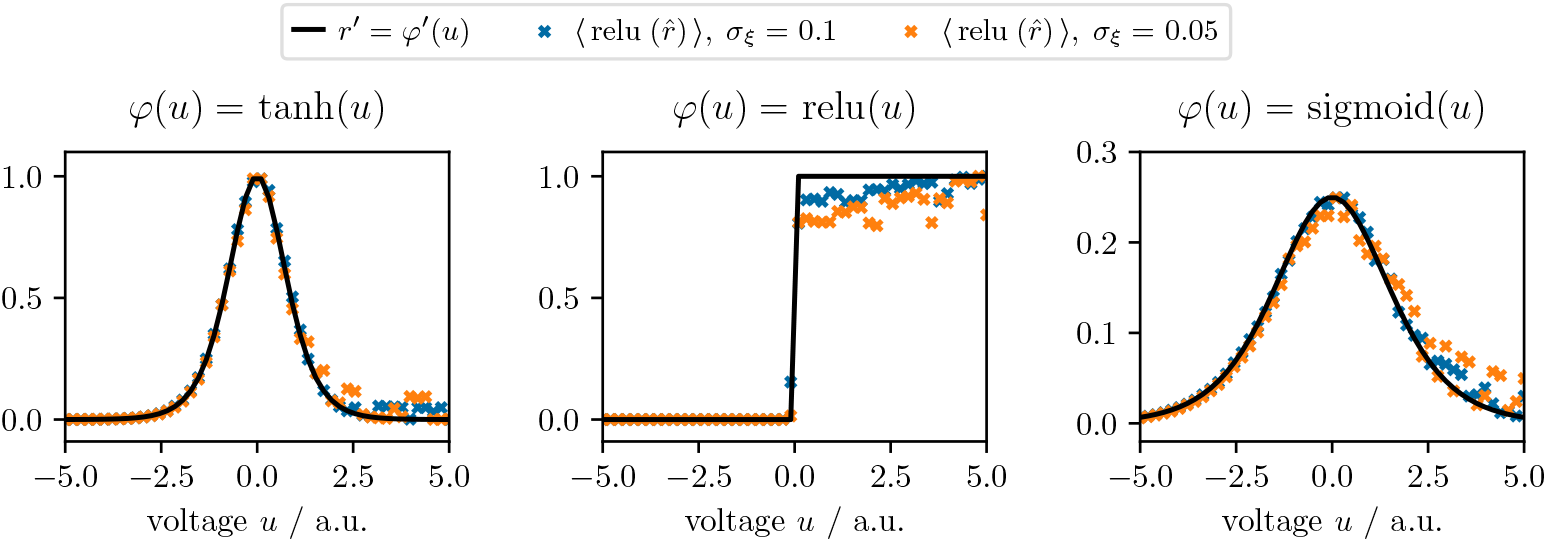
Temporal noise and non-linear dendrites enable the reconstruction of *φ*^*′*^(*u*) in error neurons. A representation unit encodes a signal voltage *u* for *T*_pres_ = 100 dt, and white noise *ξ* ∼ 𝒩(0, *σ*_*ξ*_) is sampled at every dt. After high-pass filtering with *τ*_*ξ*_ = 10 dt, the mean dendritic activity accurately tracks *φ*^*′*^(*u*) for different activation functions. The overall normalization of each curve has been rescaled for ease of illustration.

### 6.5. Simulations

We simulated the error neuron microcircuits using the Euler forward method with discrete time steps dt, implementing the neuronal dynamics of Equations (7) to (13) and the plasticity of Equation (15). Note that neuronal and weight dynamics are applied during all time steps, without phases or information buffering. The implementation builds on the code of [67], simulating the model of dendritic error microcircuits [33], see Methods in [67] for details. In all experiments, all weight matrices are fully connected, and we allow voltages to settle during a brief settling phase of several dt. Data is fed into the networks as pairs of input and targets (supervised training), where each data point is presented for *T*_pres_ time steps. As opposed to the model of Sacramento et al., our microcircuit takes target rates *r*^tgt^ instead of voltages (Equation (12), see also below).

For parameter values, see Appendix B.

#### Cart pole and DMS task

For the cart pole task, we trained our error neuron microcircuit networks to reproduce the control properties of the linear-quadratic regulator (LQR). We randomly sampled static input vectors 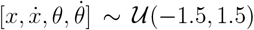 and recorded the ouput of the LQR as the teaching signal.

For validation and testing, we use a simulation of the non-linear dynamics of a cart pole, with the error neuron microcircuits acting as a controller. Note that the time step width of the cart pole simulation is equivalent to that of the microcircuits. For every time step dt, the controller receives the vector 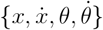 as input and produces a scalar controlling signal *F*, which is passed to the physical cart pole simulation. The cart pole dynamics are then advances by one step, closing the loop between controller and system. The cart pole is initialized with all variables 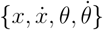 sampled uniformly from 𝒰 (−1, 1). For the angles, this corresponds to approx. *±* 57^*°*^. The cart pole is reset if |*θ*| exceeds 70^*°*^. Test performance is measured by the penalty log, i.e. how many time steps (dt = 10^−2^ ms) the cart pole is lying per epoch (20 s).

For the DMS task, we set up a network of two hidden and one output representation neurons and an equal number of error neurons. The task is constructed from train, validation and test sets of 50 stimuli pairs ({↑↑, ↑↓, ↓↑, ↓↓}) together with the corresponding target ({↑, ↓, ↓, ↑}).

The input data is a 1-dim. time series consisting of an initial fixation period of 250 ms, a first stimulus phase of length 250 ms, a delay period of random length ∼ 𝒰 (50, 250), and the second stimulus presented for 250 ms. Only at the last step of this time series, the network is provided with the target.

The encode memory at a useful timescale, we chose the conductances of the representation neurons to accommodate an effective membrane time constant of 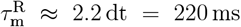. In order to propagate the effects of memory across the rate level to downstream areas, we use retrospective representation neurons [40], for which 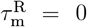. Error neurons are quasi-instantaneous, 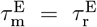. For the activation function of output neurons, we picked 20 × tanh(*x −* 0.5), representing a smooth binary classification function.

#### Teacher-student experiments

The experiments in Fig. 3 are designed to enable a fair comparison with the models of [33] and [25]. To do so, we generate a synthetic dataset by feeding random inputs to a teacher ANN with randomly initialized, fixed weights, and recording the output of the teacher network. Inputs and weights are sampled such that the input-output mapping is non-trivial (that is, non-linear).

To demonstrate that our model scales to train many cortical areas, we increase the task difficulty by scaling the architecture of the teacher and student networks from [2-1] neurons (no hidden layer) to [4-2-1], [8-4-2-1], [16-8-4-2-1] and finally [32-16-8-4-2-1] neurons (3 hidden layers). Neuron numbers double in order to make training of all areas necessary, such that errors actually need to be backpropagated, instead of a relying on learning only of output area weights. The dataset is then fed into our network in the classification configuration, such that representations propagate downstream to match the teacher output.

We also train the model of Sacramento et al. [33], using the implementation described in [67]. To enable a fair comparison, we augment the model with the prospective coding mechanism, and initialize the model with ideal lateral weights 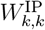 and 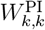 (‘self-predicting state’). In order to facilitate tight matching of lateral inhibition, we set the learning rates of 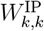 to be twice that of the efferent feed-forward weights 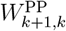. The top-down weights 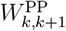 connecting pyramidal cells are adjusted in the same way as in our model, setting them to the transpose of forward-weights 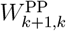 plus noise (Section 6.3). The matching lateral weights are set to the noiseless (!) transpose after every time step, i.e.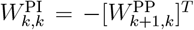. We include this ideal weight transfer in order to more objectively represent the computational power of the model by Sacramento et al.

Furthermore, their model takes voltages 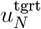 as targets; in order to compare all models on the same task, we thus feed the teacher output *r*^tgt^ into the model as the voltage 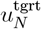. For a fair comparison, the MSE loss is then calculated based on the somatic prospective 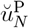 in the output area instead of the rate. Note that it is still the rate 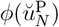 which is propagated to lower areas.

Finally, we also train the model of Mikulasch et al. [25] in the following configuration: inputs are presented to the top-most area, and top-down predictions are propagated to the lowest level, which receives the target *r*^tgt^. We have implemented two non-linear variants of dendritic hPC, but observe no difference in performance; we thus have included the linear model in Fig. 3. As described in [25], the computations performed in the model by Mikulasch et al. and that of Sacramento et al. are related. We relate them in detail in Appendix C. Compared to the model of Sacramento et al., our implementation of dendritic hPC abstracts away the interneurons, by setting the interneuron voltage to the rate of the lateral pyramidal cells, 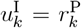; like the representation units, all interneurons have linear activation functions. In all other details, our simulations of dendritic hPC follow the same steps as described above for the model by Sacramento et al., see Appendix C.

To compute the angle between weight updates ∠(Δ*W*, Δ*W* ^ANN^), we computed Δ*W* for each model (ANN, error neuron microcircuit, Sacramento et al., dendritic hPC) for each input/target pair of the test set, and calculate the cosine similarity between Δ*W* ^ANN^ and the other models. In order to compare the error neuron microcircuit with skip connections to the hierarchical ANN, we only include the Δ*W* of neighboring areas in the angle calculation.

#### Image generation tasks

For the generative tasks, we selected one example image per digit class from the MNIST dataset, and trained the network on the pairs of classes/pixels. Image generation is fully deterministic; each class always produces the same image. There are no separate train and test sets in this task.

To model neuron variability (Fig. 5 a), we initialized each neuron individually with an activation function with slope ∼ 𝒩(1, *σ*_act_) and offset *∼* 𝒩(0, *σ*_act_).

## Acknowledgments

This work is part of the project “Putting ‘Backpropagation and the brain’ to the test”, funding KM under the Postdoc.Mobility program of the Swiss National Science Foundation. Part of this work has also been funded by the Volkswagen Foundation under the call “NEXT — Neuromorphic Computing”. KM gratefully acknowledges support by both initiatives.

We would like to thank Laura Kriener, Kenji Doya, Fabian Mikulasch, Kota Shirahata and Lucy Palmer for fruitful discussions, as well as the high performance computing support at OIST and the University of Bern.

## Author contributions

Model development: KM, together with MAP, AG and KAW. Simulations: KM and IJ. Writing: KM. Review & editing: all authors.

## Appendices

### A. Additional simulation results

**Figure A1:**
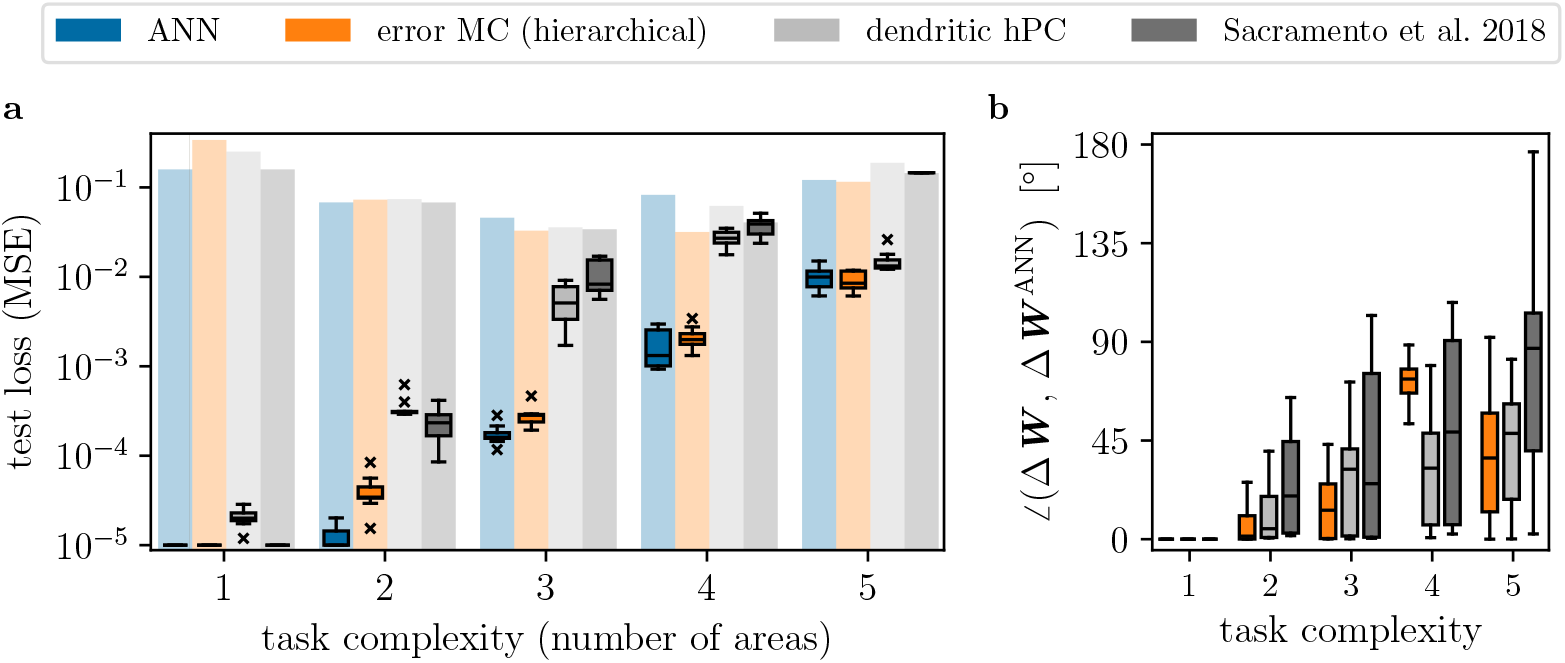
error neuron microcircuit performs equally well with realistic *and* strictly hierarchical connectivity. As opposed to dendritic hPC and Sacramento et al. [33], our model implements skip connectivity. To demonstrate that the difference in performance does not stem from the different architectures, we repeat the experiment of Fig. 3 with strictly hierarchical connectivity for the error neuron microcircuits. Note that performance is essentially unaffected, while the weight updates are much closer to those of an ANN.

**Figure A2:**
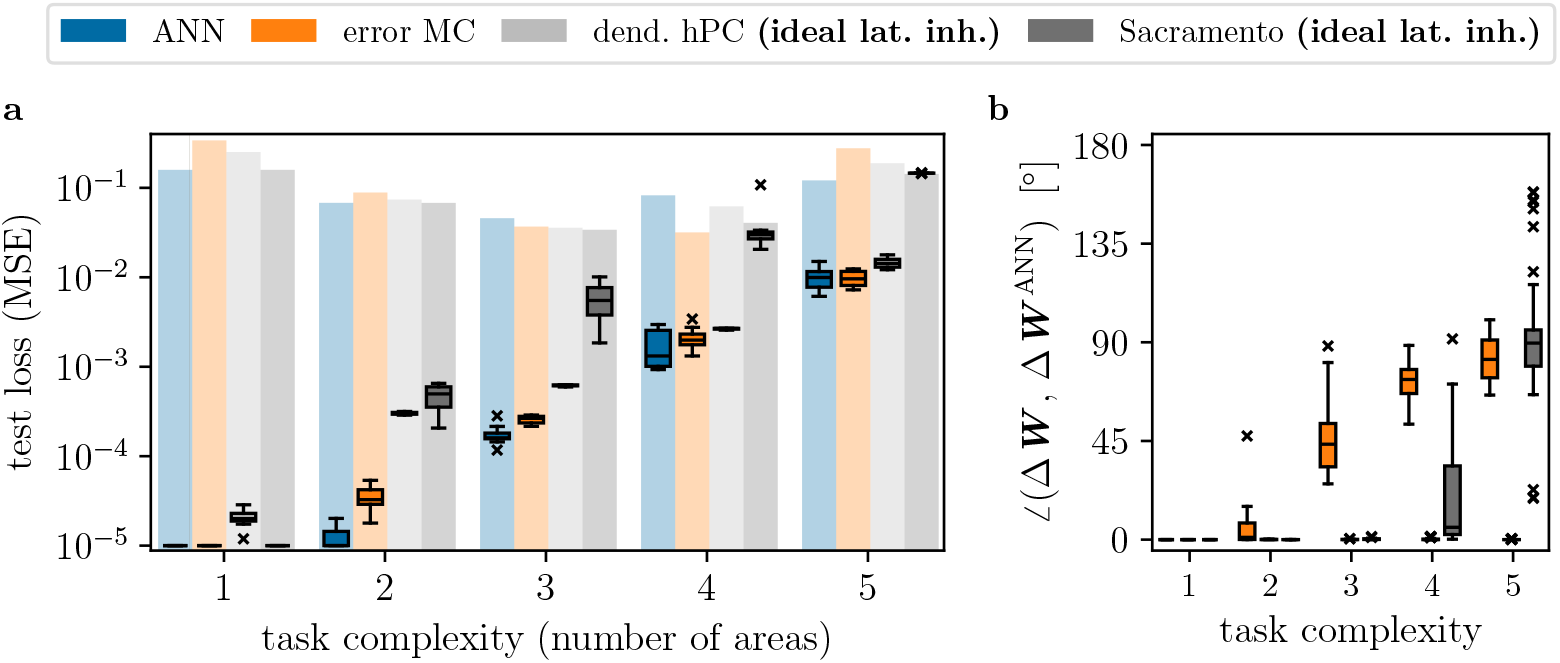
Dendritic error construction requires ideal lateral inhibition. Models where errors are constructed on dendrites (instead of explicit error neurons) require tight balance between top-down prediction and lateral inhibition. In Fig. 3, we have shown that such models do not scale to many areas. We argue that one of the reasons is learning of imperfect lateral weights when training multiple areas. To demonstrate this, we repeat the experiment with lateral inhibition set to ideal weights during learning. We observe that dendritic hPC performs much better under such artificial weight copying, but not on par with error neuron microcircuits (without weight transfer) or the ANN.

**Figure A3:**
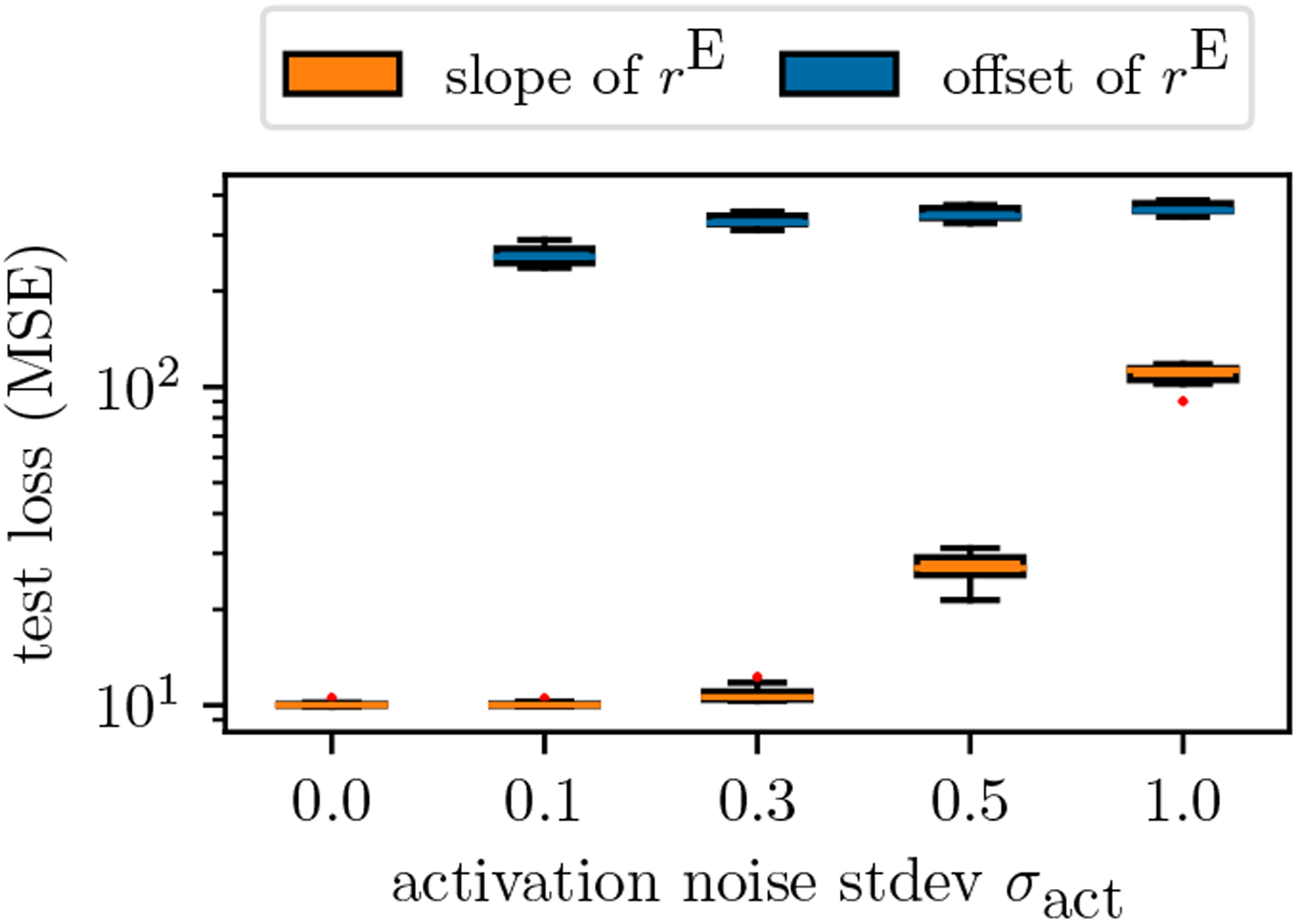
Network performance is highly sensitive to offsets of error signals. We repeat the experiment of Fig. 5 a, simulating noise only on the activation functions of error neurons, either on the slope or offset. Under slope noise, performance is stable, with similar results to offset and slope noise on representation units (Fig. 5 a). However, any noise on the offset immediately hinders learning, demonstrating the dependence of the network on useful error signals.

### B. Simulations parameters

For all simulations, we set the resting potential to *E*_l_ = 0. We set the capacitance as *C*_m_ = 1, thus conductances have units ms^−1^. All networks (error neuron microcircuit, ANN, Sacramento et al. and dendritic hPC) are trained with vanilla gradient descent with batch size 1. All simulations were performed with 10 different random seeds.

**Table B1:**
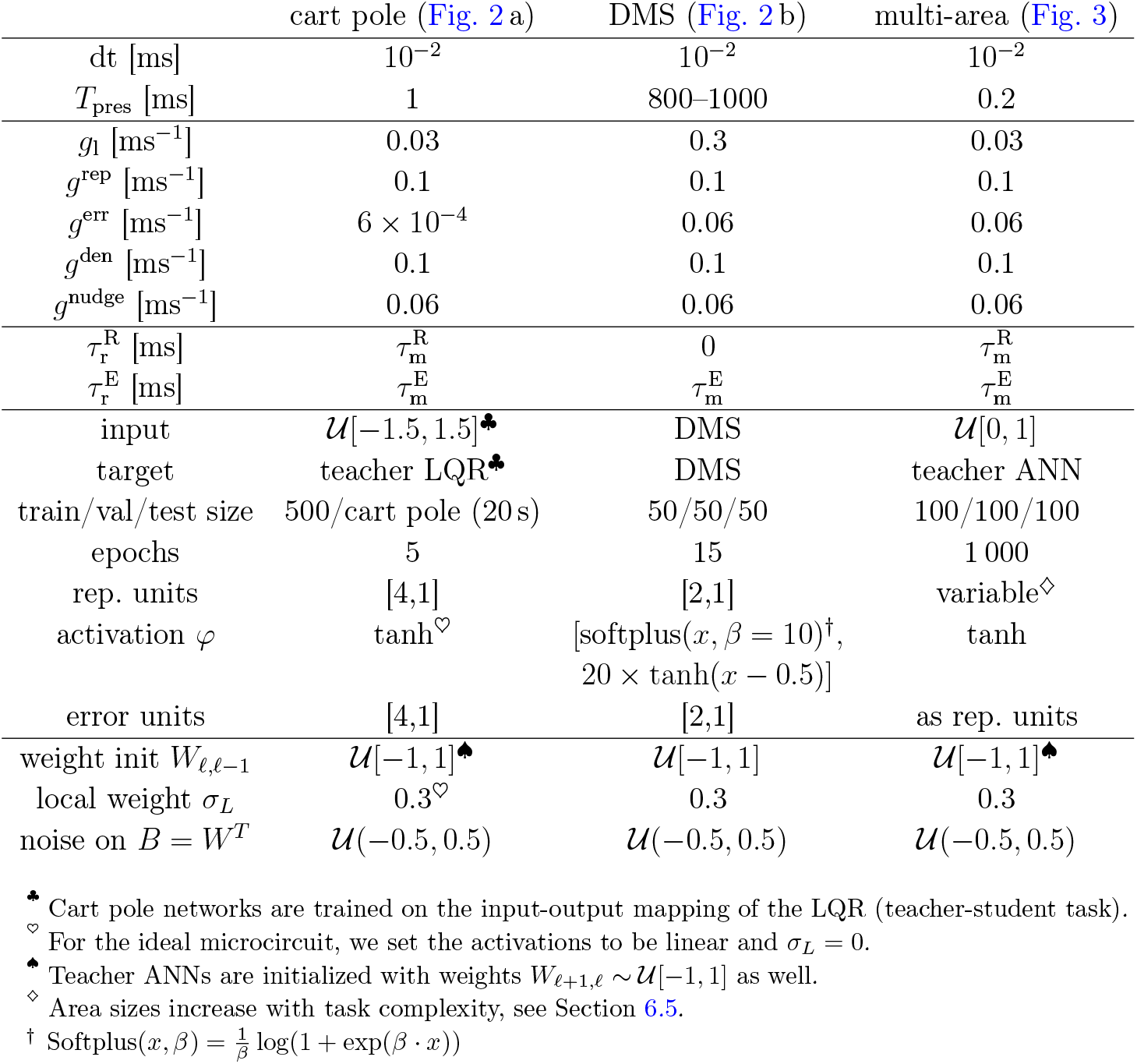
Parameters for microcircuit model simulations.

|  | cart pole (Fig. 2 a) | DMS (Fig. 2 b) | multi-area (Fig. 3) |
| --- | --- | --- | --- |
| dt [ms] | $10^{-2}$ | $10^{-2}$ | $10^{-2}$ |
| $T_{\text{pres}}$ [ms] | 1 | 800–1000 | 0.2 |
| $g_1$ [$\text{ms}^{-1}$] | 0.03 | 0.3 | 0.03 |
| $g^{\text{rep}}$ [$\text{ms}^{-1}$] | 0.1 | 0.1 | 0.1 |
| $g^{\text{err}}$ [$\text{ms}^{-1}$] | $6 \times 10^{-4}$ | 0.06 | 0.06 |
| $g^{\text{den}}$ [$\text{ms}^{-1}$] | 0.1 | 0.1 | 0.1 |
| $g^{\text{nudge}}$ [$\text{ms}^{-1}$] | 0.06 | 0.06 | 0.06 |
| $\tau_r^{\text{R}}$ [ms] | $\tau_m^{\text{R}}$ | 0 | $\tau_m^{\text{R}}$ |
| $\tau_r^{\text{E}}$ [ms] | $\tau_m^{\text{E}}$ | $\tau_m^{\text{E}}$ | $\tau_m^{\text{E}}$ |
| input | $\mathcal{U}[-1.5, 1.5]^{\clubsuit}$ | DMS | $\mathcal{U}[0, 1]$ |
| target | teacher LQR$\clubsuit$ | DMS | teacher ANN |
| train/val/test size | 500/cart pole (20 s) | 50/50/50 | 100/100/100 |
| epochs | 5 | 15 | 1 000 |
| rep. units | [4,1] | [2,1] | variable$\diamond$ |
| activation $\varphi$ | $\tanh^{\heartsuit}$ | $[\text{softplus}(x, \beta = 10)^{\dagger}, 20 \times \tanh(x - 0.5)]$ | $\tanh$ |
| error units | [4,1] | [2,1] | as rep. units |
| weight init $W_{\ell, \ell-1}$ | $\mathcal{U}[-1, 1]^{\spadesuit}$ | $\mathcal{U}[-1, 1]$ | $\mathcal{U}[-1, 1]^{\spadesuit}$ |
| local weight $\sigma_L$ | $0.3^{\heartsuit}$ | 0.3 | 0.3 |
| noise on $B = W^T$ | $\mathcal{U}(-0.5, 0.5)$ | $\mathcal{U}(-0.5, 0.5)$ | $\mathcal{U}(-0.5, 0.5)$ |

$\clubsuit$ Cart pole networks are trained on the input-output mapping of the LQR (teacher-student task).
$\heartsuit$ For the ideal microcircuit, we set the activations to be linear and $\sigma_L = 0$ .
$\spadesuit$ Teacher ANNs are initialized with weights $W_{\ell+1, \ell} \sim \mathcal{U}[-1, 1]$ as well.
$\diamond$ Area sizes increase with task complexity, see Section 6.5.
† $\text{Softplus}(x, \beta) = \frac{1}{\beta} \log(1 + \exp(\beta \cdot x))$

**Table B2:**
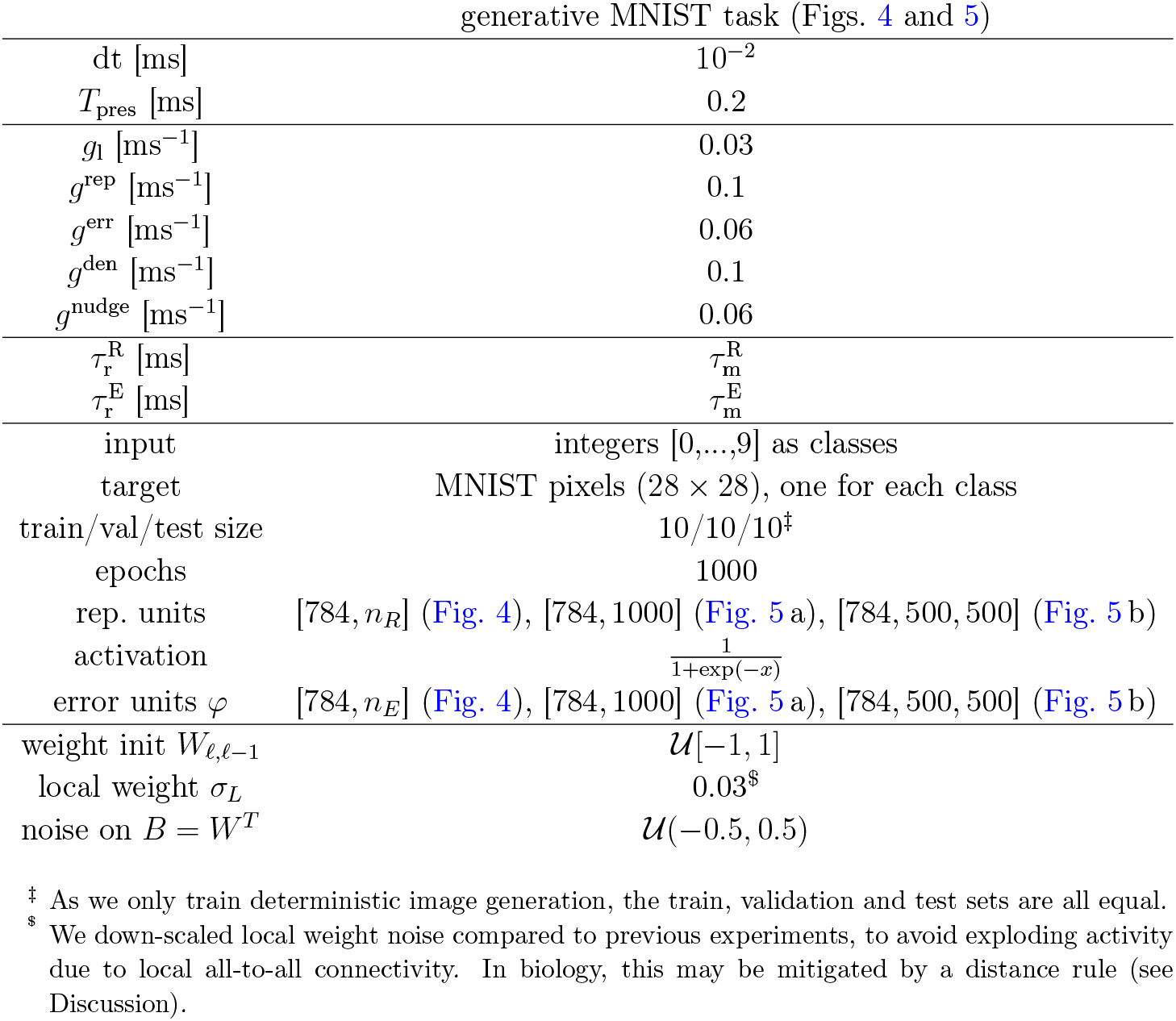
Parameters for microcircuit model simulations (cont’d).

### C. Implementation of dendritic hierarchical PC

We have adapted the model by Mikulasch et al. [25] in the following way. As explained in 2 of [25], dendritic hPC can be related to the model of Sacramento et al. [33] by inverting the hierarchy of cortical areas. This amounts to a generative and classifying configuration, respectively (see Fig. 1). As we are comparing the networks on an arbitrary, deterministic input-output mapping task (cf. Fig. 3), there is no actual distinction between generative and classifying configurations; i.e., labels and targets are defined arbitrarily and can be exchanged. To allow for a fair comparison across all models, we therefore swap input and target for dendritic hPC. This effectively trains the network to solve the same task as all other models, which are run in their classification configuration. For example, for 3 areas, all networks receive 8 inputs and process them using three areas with [4, 2, 1] neurons.

We implement dendritic hPC following equations VIII to XIII in Box 2 of [25]. By exchanging basal and apical compartment labels and rewriting the soma dynamics, one can see that dendritic hPC is in fact largely computationally equivalent to the model by Sacramento et al.: Starting from the dendritic microcircuit model [33], we can set 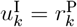, drop the weight 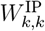 from the theory, and equate the swapped compartments, 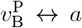 and 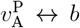. Soma and dendrite dynamics can be seen to be equivalent by redefining 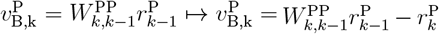. Conductances can be chosen to mirror the dynamics of Eqs. VIII, IX and XI. Weights are then related by 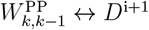 (Eq. XIII), 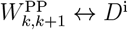 (Eq. XII), and 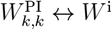 (Eq. X).

To establish a fair comparison, we further make the following changes:

- we set both compartments to be instantaneous, *τ*_b_ = *τ*_a_ = 0, equal to the other models in this work.
- we implement the same approximation of PAL as in the other models by setting 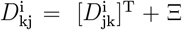, with Ξ fixed noise. This means that we do not implement Eq. XII. It also greatly reduces computational complexity, as the index k in 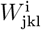 can be dropped, reducing the tensor to a matrix.
- the theory of [25] is purely linear. To introduce non-linear activations *φ*, we re-label the variable *r* (Eq. XI) to *u*, representing the soma, and *r* now representing the output rate.
- to mitigate the relaxation problem, we augment neuron outputs with the prospective coding mechanism, 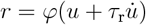, as in the other models.
- to account for the non-linear activation functions, we optionally implement the Urbanczik-Senn learning rule in the form 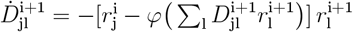.
- we use the same conductances *{g*_l_, *g*^rep^, *g*^err^} as in the other models (for the model of Sacramento et al., *g*^rep^ = *g*^bas^ and *g*^err^ = *g*^api^). This can be interpreted as a generalized parametrization of the precision weighting (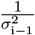 and 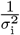 in Eq. XI of [25]) of both compartments.

To facilitate tight balance, we set the learning rate of 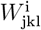 (self-inhibition in area i) to be double that of 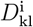 (efferent weights of area i, projecting to areas i − 1 and i + 1). Furthermore, we initialize each network with ideal lateral weights, 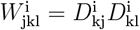. We compare the different implementations of dendritic hPC in Fig. C1.

**Figure C1.**
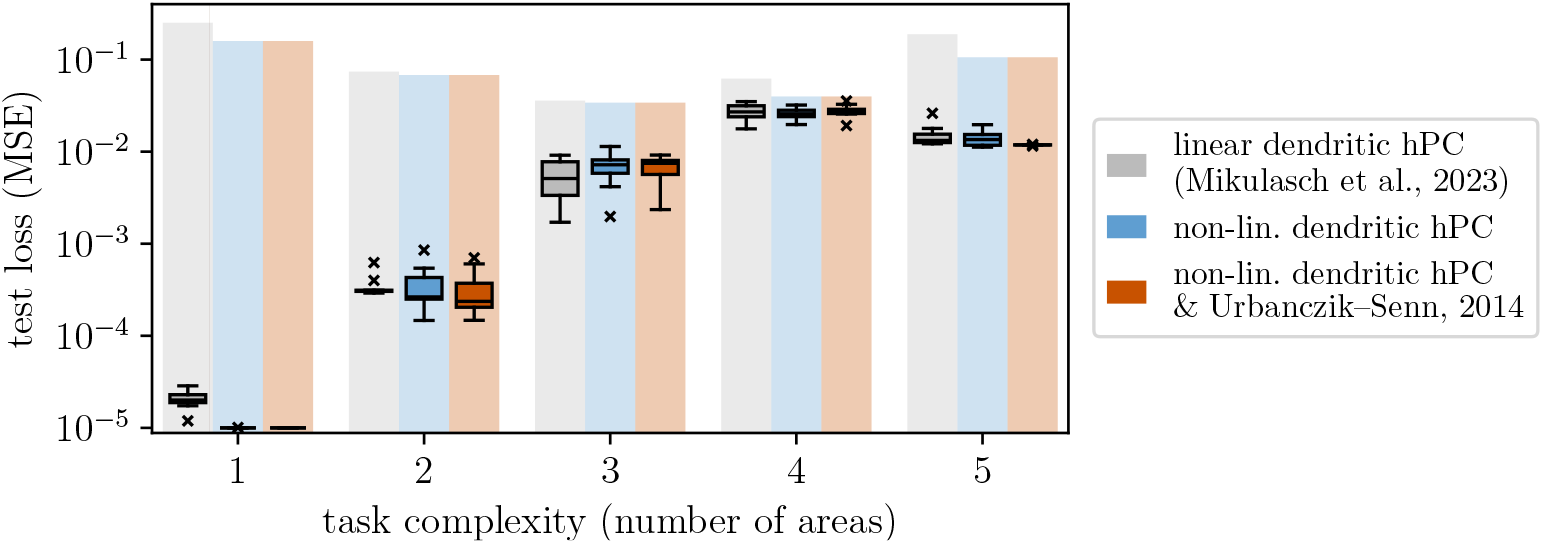
Three different implementations of dendritic hPC for learning non-linear tasks. The original model of dendritic hPC by Mikulasch et al. [25] is a purely linear theory. For a fair comparison to our model on a non-trivial benchmark, we implement the original model (grey) and two variants, where linear activations are replaced by non-linearities (light blue), and additionally, the learning rule for 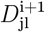 is adapted to the Urbanczik-Senn rule [133] (brown). As both modifications do not significantly change performance, we have used the original (grey) model in Fig. 3.

### D. Alternative description of our model with top-down errors in representation dendrites

As discussed in Section 4, particularly Fig. 6, our model admits an alternative parametrization, where representation dendrites encode top-down errors instead of predictions. Here, we show the mathematical equivalence of both parametrizations.

To reformulate our neuron dynamics as in dendritic hPC (Box 2 in [25]), we need to modify the somatic and representation dendrite dynamics of representation units:

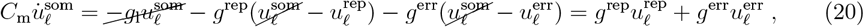

and adding a term to 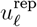,

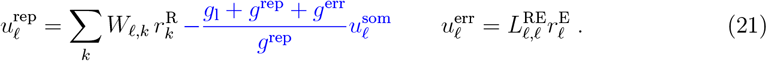

I.e., the representation compartment now compares the conductance-weighted somatic voltage to top-down predictions. By plugging Equation (21) into Equation (20), one can see that 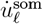 remains unchanged.

We adapt the learning rule of Equation (4) by using the top-down prediction in place of 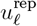,

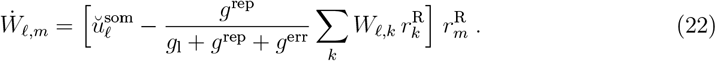

